# Evolutionary Origins of Self-sustained Kai Protein Circadian Oscillators

**DOI:** 10.1101/2024.07.23.604570

**Authors:** Atsushi Mukaiyama, Yoshihiko Furuike, Kumiko Ito-Miwa, Yasuhiro Onoue, Kota Horiuchi, Eiki Yamashita, Shuji Akiyama

## Abstract

Light–dark cycles affect photosynthetic efficiency in autotrophic cyanobacteria; therefore, determining whether ancient cyanobacteria possessed a self-sustained circadian clock when oxygenic photosynthetic systems were established is an important issue in chronobiology. Here we examined the oscillation of the clock protein KaiC in modern cyanobacteria, as well as the function and structure of ancestral Kai proteins, to determine the evolutionary origin of the self-sustained Kai-protein oscillators. The results show that the oldest double-domain KaiC in ancestral bacteria lacked the factors functionally and structurally essential for rhythmicity. The ancestral Kai proteins had acquired these factors through molecular evolution that occurred around Global Oxidation and Snowball Earth events, and were eventually inherited as a self-sustained circadian oscillator by the most recent common ancestor of cyanobacteria capable of oxygenic photosynthesis. This autonomous Kai protein oscillator was further inherited by most freshwater and marine cyanobacteria present today as an autotrophic basis for time-optimal acquisition and consumption of energy from oxygenic photosynthesis.

## Main

The most recent common ancestor (MRCA) of cyanobacteria emerged approximately 3–2 billion years (Ga) ago ^1-6^ and evolved into the current ecosystem through the Great Oxidation event (GOE) 2.3 Ga ago ^7^, at least two Snowball Earth events (SEEs) 2.4 and 0.7 Ga ago ^8-10^, and the Neoproterozoic Oxygenation event (NOE) 0.8–0.6 Ga ago ^11,12^. Studies of fossils and molecular evolution models suggest that the MRCA of cyanobacteria already possessed primitive oxygenic photosynthetic systems ^1,2,4^. Because the efficiency of photosynthesis is strongly influenced by light–dark environmental cycles, determining whether primitive cyanobacteria possessed a timekeeping system when photosynthesis became active in the GOE period is important to understanding the physiological origin of circadian clock systems ^13^. Primitive biological clocks are also of interest from the perspective of geoscience, and the evolution of the Earth’s rotation period from ancient times to the present remains an issue of debate ^14^.

Circadian clocks with autonomous 24-h oscillatory, temperature compensatory, and synchronous properties have been identified in various extant organisms ^15^, including bacteria ^16^, fungi ^17^, plants ^18^, and mammals ^19^. The circadian clock of cyanobacteria has been studied intensively using the *Synechococcus elongatus* PCC7942 (*Se*7942) strain as a model system ^20,21^. The clock oscillator can be reconstructed in a test tube by incubating the core protein KaiC from the *Se*7942 strain (KaiC*^Se^*) in the presence of KaiA*^Se^*, KaiB*^Se^*, and adenosine triphosphate (ATP)^22^. KaiC*^Se^* exhibits rhythms in phosphorylation ^23,24^ and interactions with KaiA*^Se^* and KaiB*^Se^* ^25-28^; however, the frequency of these rhythms is highly correlated with the ATPase activity of KaiC*^Se^* itself ^29,30^. For example, when the ATPase activity of KaiC*^Se^* doubles as a result of amino acid substitutions, the frequencies of both the *in vitro* oscillator and the intracellular rhythm also double. This process mediated by KaiC*^Se^* is known as cross-scale causality and is also observed in temperature compensation ^31^.

One hypothesis worth testing is whether primitive cyanobacteria utilized a Kai-protein oscillator. However, the concept of a protein-based oscillator is derived almost exclusively from experiments and observations using one extant species, the *Se*7942 strain. Therefore, whether Kai proteins functioned as circadian oscillators in ancestral cyanobacteria, or even in extant species other than the *Se*7942 strain, remains unknown.

## Results

### Feasibility study of reconstructing Kai-protein oscillators in extant species

Duplication of the predecessor of the *kaiC* gene and their subsequent fusion into a double-domain predecessor occurred between 3.8 and 3.5 Ga ago ^32^. We used the result of large-scale genome analyses of cyanobacteria and other species ^33^ to construct a phylogenetic tree of extant double-domain KaiC homologs (Fig. 1a and Extended Data Fig. 1). The KaiC tree included four groups: group I was composed mainly of KaiCs from freshwater cyanobacteria, including *Se*7942 (ID: 001); group II consisted of marine cyanobacteria; group III included other bacteria; and group IV was formed by other bacteria including archaea. Most freshwater cyanobacteria have only one *kaiC* of group I, but others have additional group III or IV *kaiC* originated from secondary copies or lateral transfer. Thus, these group III and IV KaiCs from freshwater cyanobacteria (gray entries in Fig. 1a) in the protein-based tree can be excluded when identifying groups I–IV and nodes based on species and growing habitant (see Supplementary Information).

**Fig. 1.**
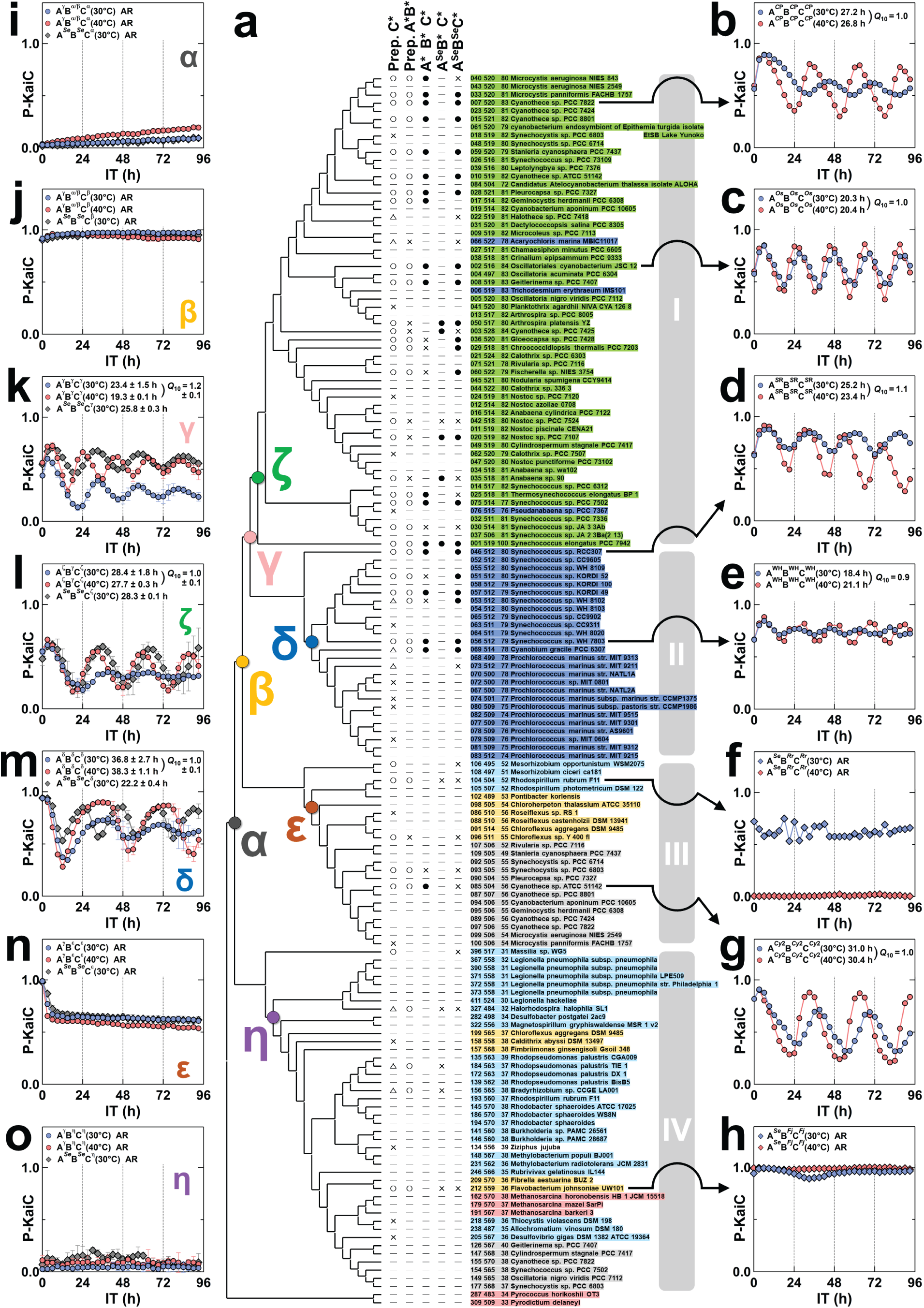
Feasibility studies of *in vitro* oscillators for extant and ancestral Kai proteins. (**a**) Phylogenetic tree of extant double-domain KaiC homologs. A detailed view of full entries is presented in Extended Data Fig. 1. Cyanobacteria in freshwater, marine cyanobacteria, proteobacteria, bacteria, and euryarchaeota/crenarchaeota are colored green, blue, cyan, yellow, and red, respectively. Group III and IV KaiCs from freshwater cyanobacteria (gray entries) are originated from secondary copies of group I *kaiC* gene or lateral gene transfer, and thus can be excluded when identifying groups I–IV and nodes based on species and growing habitant. The three numbers preceding the name of each entry indicate the ID number, the total number of amino acids, and the sequence identity with KaiC*^Se^*, respectively. A*B*C*: sets of species-specific KaiA*, KaiB*, and KaiC*. A*^Se^*B*C*: sets of KaiA*^Se^*, KaiB*, and KaiC*. A*^Se^*B*^Se^*C*: sets of KaiA*^Se^*, KaiB*^Se^*, and KaiC*. Rhythmic sets are indicated by filled circles, whereas arhythmic sets are labeled with crosses (see each result in Extended Data Fig. 3). Prep. C* and Prep. A*B* indicate whether species-specific Kai proteins were adequately expressed and purified (open circles), prepared at a level that could be analyzed (triangles), or not prepared at all (crosses). Symbols α–η represent evolutionarily important branching points for which the amino acid sequence of ancestral KaiC was restored (Fig. 2a). *In vitro* rhythm assays of A*B*C* (circles) or A*^Se^*B*C* (diamonds) at 30°C (blue) and 40°C (red) as functions of incubation time (IT) for (**b**) *Cyanothece sp.* PCC7822 (KaiC*^C^*^P^, ID: 007) in group I, (**c**) *Oscillatoriales cyanobacterium* JSC 12 (KaiC*^Os^*, ID: 002) in group I, (**d**) *Synechococcus sp.* RCC307 (KaiC*^S^*^R^, ID: 046) in group II, (**e**) *Synechococcus sp.* WH 7803 (KaiC^WH^, ID: 056) in group II, (**f**) *Rhodospirillum rubrum* F11 (KaiC*^Rr^*, ID: 104) in group III, (**g**) *Cyanothece sp.* ATCC 51142 (KaiC*^Cy^*^2^, ID: 085) in group III, and (**h**) *Flavobacterium johnsoniae* UW101 (KaiC*^Fj^*, ID: 212) in group IV. Shown are the representative examples from multiple experiments summarized in Extended Data Fig. 3. AR indicates that the period could not be determined (arrhythmic). *In vitro* rhythm assays for (**i**) A^γ^B^α/β^C^α^ (circles) and A*^Se^*B*^Se^*C^α^ (diamonds), (**j**) A^γ^B^α/β^C^β^ (circles) and A*^Se^*B*^Se^*C^β^ (diamonds), (**k**) A^γ^B^γ^C^γ^ (circles) and A*^Se^*B*^Se^*C^γ^ (diamonds), (**l**) A^ζ^B^γ^C^ζ^ (circles) and A*^Se^*B*^Se^*C^ζ^ (diamonds), (**m**) A^δ^B^δ^C^δ^ (circles) and A*^Se^*B*^Se^*C^δ^ (diamonds), (**n**) A^γ^B^ε^C^ε^ (circles) and A*^Se^*B*^Se^*C^ε^ (diamonds), and (**o**) A^γ^B^η^C^η^ (circles) and A*^Se^*B*^Se^*C^η^ (diamonds). Blue circles and gray diamonds correspond to the data taken at 30°C, and red circles correspond to the data taken at 40°C. Values are presented as means ± SD (n = 3).

*Rhodobacter sphaeroides* KaiC (KaiC*^Rs^*, ID: 194), which functions as an hourglass-like timekeeping system ^34^, and *Rhodopseudomonas palustris* TIE 1 KaiC (KaiC*^Rp^*, ID: 184), which is not self-sustained but enhances fitness in the rhythmic environment ^35^, were classified into group IV in the present KaiC tree. The overall topology and grouping of the protein-based tree of double-domain KaiCs are consistent with those of the 16S rRNA–based phylogenic tree (Extended Data Fig. 2, see Supplementary Information) as well as with a previous report ^32^.

We used the *in vitro* rhythm assay ^22^ to test the feasibility of establishing the *in vitro* oscillator for 41 representative extant species including the *Se*7942 strain at pH 8.0 and 30°C (Extended Data Fig. 3). In most KaiCs from extant species belonging to group I, circadian rhythms were observed in the presence of the corresponding species-specific KaiA and KaiB. For example, *Cyanothece sp.* PCC7822 KaiC (KaiC*^C^*^P^, ID: 007) cycled between phosphorylated and de-phosphorylated forms with a period of 27.2 h (Fig. 1b) in the presence of KaiA*^C^*^P^ and KaiB*^C^*^P^, as observed in the *Se*7942 system (Extended Data Fig. 3a). The same rhythm was observed in *Oscillatoriales cyanobacterium* JSC 12 KaiC (KaiC*^Os^*, ID: 002) (Fig. 1c). Cases in which an obvious rhythm was detected in a set (A*B*C*) corresponding to species-specific KaiA (KaiA*), KaiB*, and KaiC* are shown in filled circles in Fig. 1a. Regardless of the behaviors of the A*B*C* set, cases in which oscillation was detected when KaiA* and KaiB* were replaced by KaiA*^Se^* and KaiB*^Se^*, respectively (A*^Se^*B*^Se^*C*), are also marked with filled circles in a different column (Fig. 1a). In approximately 80% of group I A*B*C* trials, the rhythm showed a period of 25.1 ± 5.0 h at 30°C, whereas in approximately 70% of the examined group I A*^Se^*B*^Se^*C* sets, the period was 23.4 ± 4.5 h (Extended Data Fig. 3a).

In species with KaiCs classified into groups II–IV that did not have a homologous gene for *kaiA*, KaiA*^Se^* was hypothetically used for the *in vitro* rhythm assay (A*^Se^*B*C*). Robust circadian rhythms were observed for most of the examined A*B*C* sets in the first half of the group II entries (67%, Extended Data Fig. 3b), such as *Synechococcus sp.* RCC307 (KaiC*^S^*^R^, ID: 046) (Fig. 1d) and *Synechococcus sp.* WH 7803 (KaiC^WH^, ID: 056) (Fig. 1e). In the other half of group II, which consisted of *Prochlorococcus* strains lacking species-specific KaiA, most KaiCs could not be expressed or purified. We were able to purify KaiC from *Prochlorococcus marinus str.* MIT 9211 (ID: 073); however, it showed no rhythm, even in the corresponding A*^Se^*B*^Se^*C* set (Fig. 1a and Extended Data Fig. 3b).

The rhythmic fractions of the group III and IV entries were relatively low (Fig. 1a): 50% and 0% for the A*B*C* and A*^Se^*B*^Se^*C* sets of group III, respectively (Extended Data Fig. 3c), and 0% for both the A*^Se^*B*C* and A*^Se^*B*^Se^*C* sets of group IV (Extended Data Fig. 3d). For example, the KaiCs from *Rhodospirillum rubrum* F11 (KaiC*^Rr^*, ID: 104) in group III (Fig. 1f) and *Flavobacterium johnsoniae* UW101 (KaiC*^Fj^*, ID: 212) in group IV (Fig. 1h) did not show self-sustained oscillation in their A*^Se^*B*C* and A*^Se^*B*^Se^*C* sets. Group-III *Cyanothece sp.* ATCC 51142 (KaiC*^Cy^*^2^, ID: 085) was the only example of self-sustained oscillation in the A*B*C* set examined for the group III and IV (Fig. 1g).

These results suggest that not all extant KaiC homologs with a double-domain architecture have a circadian clock function, which raises the question of how and when KaiC acquired its function as a self-sustained circadian clock.

### *In vitro* reconstruction of ancestral Kai protein oscillators

We restored the possible amino acid sequences of ancestral KaiCs corresponding to seven evolutionary branching points (α–η) (Fig. 1a and Extended Data Fig. 1, see Methods). Ancestral KaiCs consist of 509–560 amino acids and contain an N-terminal CI domain and a C-terminal CII domain (Fig. 2a). Similar to KaiC*^Se^*, all ancestral KaiCs retain catalytic glutamates essential for ATPase activity in the CI domain and kinase/phosphatase activity in the CII domain, as well as a potential dual phosphorylation site (S431 and T432 in KaiC*^Se^*) in the CII domain (Fig. 2a). A phylogenetic tree of extant KaiB homologs was constructed (Extended Data Fig. 4) and compared with that of the double-domain KaiC homologs (Fig. 1a). According to the similarities in the taxonomic patterns of the belonging species (see coloring by growing habitat), the branches corresponding to the γ, δ, ε, and η points were assigned to the KaiB tree (Extended Data Fig. 4), whereas α and β were identified as a point (α/β) with no clear distinction between them. Consistent with a previous study showing that ancestral KaiA appeared at a later time than ancestral KaiB and KaiC ^32,36^, we were only able to assign three later branches for the γ, δ, and ζ points in our tree of double-domain KaiA homologs (Extended Data Fig. 5). Because ancient seawater was neutral (pH ∼ 7) ^37^, whereas the current seawater is slightly basic (pH ∼ 8), and because some cyanobacteria undergo a change in intracellular pH in response to the pH of their environment ^38^, these ancestral forms of Kai proteins were expressed and purified for an *in vitro* rhythm assay at pH 7.0 and a temperature of 30°C (see Supplementary Information).

**Fig. 2.**
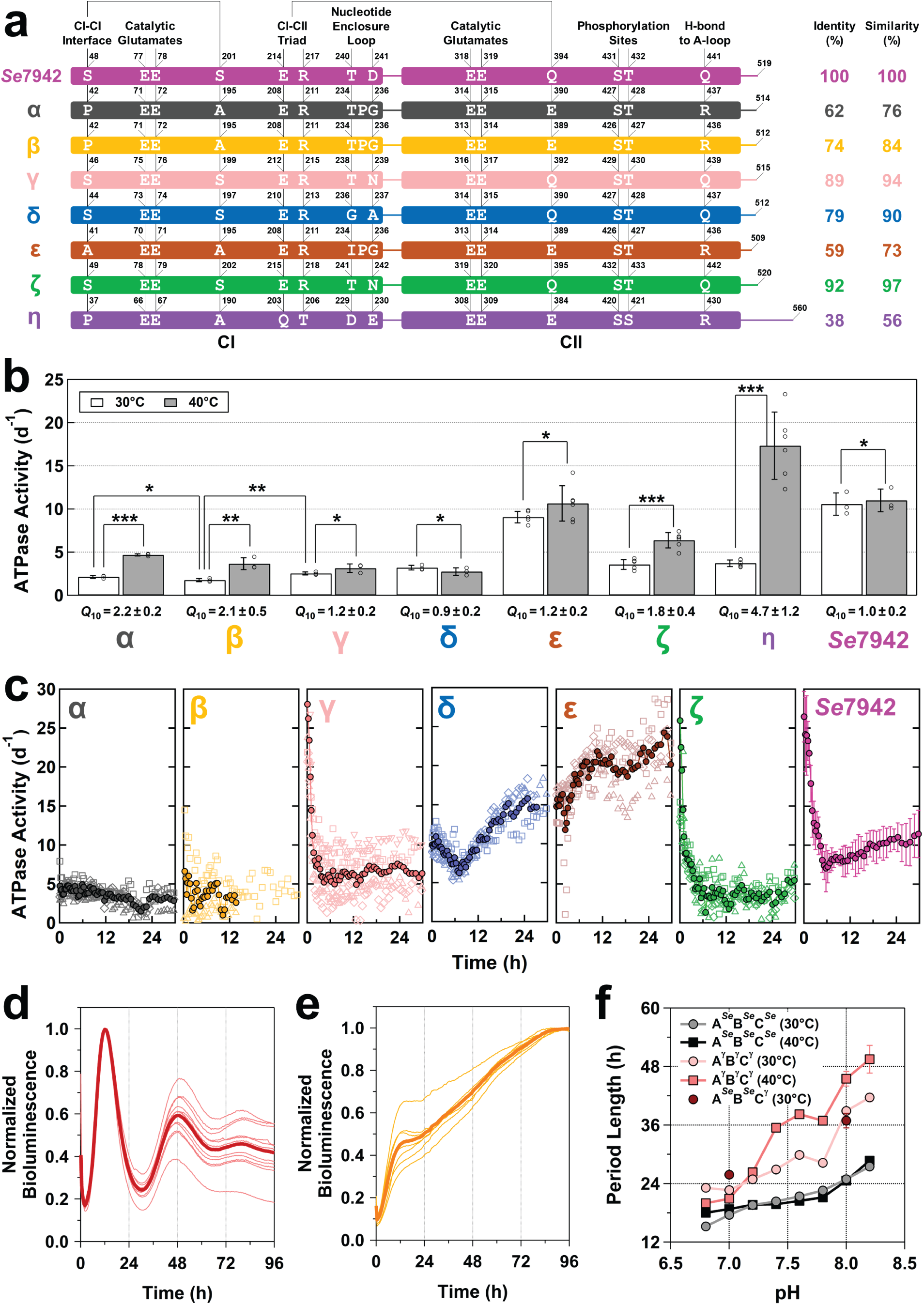
Domain architecture and biochemical properties of ancestral KaiCs. (**a**) Key residues and motifs mapped to the domain compositions of KaiC*^Se^*, KaiC^α^, KaiC^β^, KaiC^γ^, KaiC^δ^, KaiC^ε^, KaiC^ζ^, and KaiC^η^. (**b**) Temperature dependence of the steady-state ATPase activity of ancestral KaiCs and KaiC*^Se^*. Open circles correspond to raw data shown in Extended Data Table. 1. The ATPase activities at 30°C and 40°C were analyzed using *t* test; *Insignificant (P ≥ 0.05), **significant (P < 0.05), and ***significant (P < 0.005). The ATPase activities of KaiC^α^, KaiC^β^, and KaiC^γ^ at 30°C were analyzed using one-way ANOVA (F_2,6_ = 5.143, P = 0.004752) and the Bonferroni post hoc t test; *Insignificant (P ≥ 0.05/3), **significant (P < 0.05/3), and ***significant (P < 0.005/3). (**c**) Pre–steady-state analysis of the ATPase activity of ancestral KaiCs and KaiC*^Se^*. Dark-colored circles for ancestral KaiCs correspond to the mean from independent measurements (open squares, triangles, and diamonds). The data of KaiC*^Se^* were taken from the previous report (n = 3) ^29^. (**d, e**) Bioluminescent rhythm assays for *Se*7942 reporter strains carrying *kaiA^Se^*: *kaiB^Se^*: *kaiC*^γ^ and *kaiA^Se^*: *kaiB^Se^*: *kaiC*^β^, respectively, but without *kaiC^Se^*, were performed under continuous-light illumination condition at 30°C. Thin and thick lines represent individual (n = 12 and 7 for panels **d** and **e**, respectively) and average traces, respectively. The period length (32.9 ± 0.7 h) of the strain with *kaiA^Se^*: *kaiB^Se^*: *kaiC*^γ^ was consistent with that of the *in vitro* A^γ^B^γ^C^γ^ set at pH 7.6–7.8 at 30°C (panel **f**). (**f**) pH-dependence of the period length for the *in vitro* A^γ^B^γ^C^γ^ and A*^Se^*B*^Se^*C*^Se^* sets at 30°C and 40°C. The error bars represent the propagation of errors in fitting the sine function to the observed rhythmic data. The data for the A*^Se^*B*^Se^*C^γ^ set at 30°C at pH 7.0 and 8.0 were presented by brown circles as means ± SD (n = 3).

A hypothetical oldest set (A^γ^B^α/β^C^α^) consisting of KaiA^γ^, KaiB^α/β^, and KaiC^α^ did not show any signs of oscillation, and this was also observed in the A*^Se^*B*^Se^*C^α^ set (Fig. 1i). Similar results were obtained for the second oldest A^γ^B^α/β^C^β^ and A*^Se^*B*^Se^*C^β^ sets (Fig. 1j). By contrast, the A^γ^B^γ^C^γ^ and A*^Se^*B*^Se^*C^γ^ sets exhibited robust oscillations with periods of 23.4 ± 1.5 and 25.8 ± 0.3 h (Fig. 1k), respectively. Similarly, both the ζ sets (A^ζ^B^γ^C^ζ^: 24.8 ± 3.2 h, A*^Se^*B*^Se^*C^ζ^: 28.3 ± 0.1 h, Fig. 1l) and the δ sets (A^δ^B^δ^C^δ^: 36.8 ± 2.7 h, A*^Se^*B*^Se^*C^δ^: 22.2 ± 0.4 h, Fig .1m) showed rhythmic oscillation. Neither the ε sets nor the η sets (A^γ^B^ε^C^ε^, A*^Se^*B*^Se^*C^ε^, A^γ^B^η^C^η^, nor A*^Se^*B*^Se^*C^η^) exhibited any rhythmicity (Fig. 1n and 1o).

These results indicate that the oldest KaiC^α^ (62% identity to KaiC*^Se^*, Fig. 2a) and the subsequent KaiC^β^ (74% identity) lacked oscillatory ability and that this ability was acquired during the transition from KaiC^β^ to KaiC^γ^ (89% identity). Consistent with this interpretation, the ATPase activities of KaiC^α^ and KaiC^β^ were temperature-dependent (*Q*_10_: 2.1–2.2), whereas those of KaiC^γ^ and KaiC^δ^ were temperature-compensated (*Q*_10_: 0.9–1.2) (Fig. 2b and Extended Data Table. 1). Pre–steady-state analysis of the ATPase activity supported this interpretation: KaiC*^Se^*exhibits damped oscillation of ATPase activity even in the absence KaiA*^Se^* and KaiB*^Se^* ^29^, and similar characteristics were observed in KaiC^γ^, KaiC^δ^, and KaiC^ζ^ (Fig. 2c). In addition, an *Se*7942 reporter strain in which *kaiC^Se^* was replaced by *kaiC*^γ^ showed a bioluminescent rhythm under constant-light illumination conditions at 30°C (Fig. 2d and 2e), and its period length (32.9 ± 0.7 h) was consistent with that of the *in vitro* A^γ^B^γ^C^γ^ set at pH 7.6–7.8 and 30°C (Fig. 2f). The acquisition of oscillatory abilities that occurred in the transformation from KaiC^β^ to KaiC^γ^ is not a wild hypothesis confined to the test tube but a possible evolutionary event that could have occurred in primitive cyanobacteria.

We found that the period length of the A^γ^B^γ^C^γ^ set was sensitive to ambient pH. The cycle length, which was 39 h at pH 8.0 and 30°C, became shorter under acidic conditions, finally stabilizing at 23.4 ± 1.5 h at pH 7.0 and 30°C (Fig. 2f). The A*^Se^*B*^Se^*C*^Se^*set showed a similar tendency, although it was less pH-dependent than the A^γ^B^γ^C^γ^ set. The results for the A*^Se^*B*^Se^*C^γ^ set (Fig. 1k and 2f) suggest that the pronounced pH sensitivity of the period length is derived from KaiC^γ^ itself. Since the intracellular pH of cyanobacteria changes in response to the external pH ^38^ and that the pH of ancient seawater was ∼7 ^37^, the period length at pH 7.0 and 30°C is reasonably concluded to be a better estimate for ancestral types than that at pH 8.0 and 30°C.

### Structural evolution of ancestral KaiC

To explore the structural origin of the autonomous oscillatory properties acquired in the transition from β to γ, we solved the X-ray crystal structures for KaiC^β^, KaiC^γ^, and KaiC^δ^ (Extended Data Table. 2). The overall structures of KaiC^γ^ and KaiC^δ^ were similar to that of KaiC*^Se^*(Fig. 3a), and their main chains could be superimposed with a root-mean-squared deviation (RMSD) of 1.1 Å. The similarity between KaiC^γ^ and KaiC*^Se^* was confirmed by the superimposition of other parts using one particular CII domain (star in Fig. 3b). By contrast, the structure of KaiC^β^ differed markedly from that of KaiC^γ^ (main-chain RMSD of 2.0 Å). When KaiC^β^ and KaiC^γ^ were superimposed using one of the CII domains (star in Fig. 3c), the other CI and CII domains did not overlap (main-chain RMSD of 5.3 Å). These findings suggest that ancestral KaiCs acquired the basic architecture for establishing the autonomous oscillator found in KaiC*^Se^* during the transition from KaiC^β^ to KaiC^γ^.

**Fig. 3.**
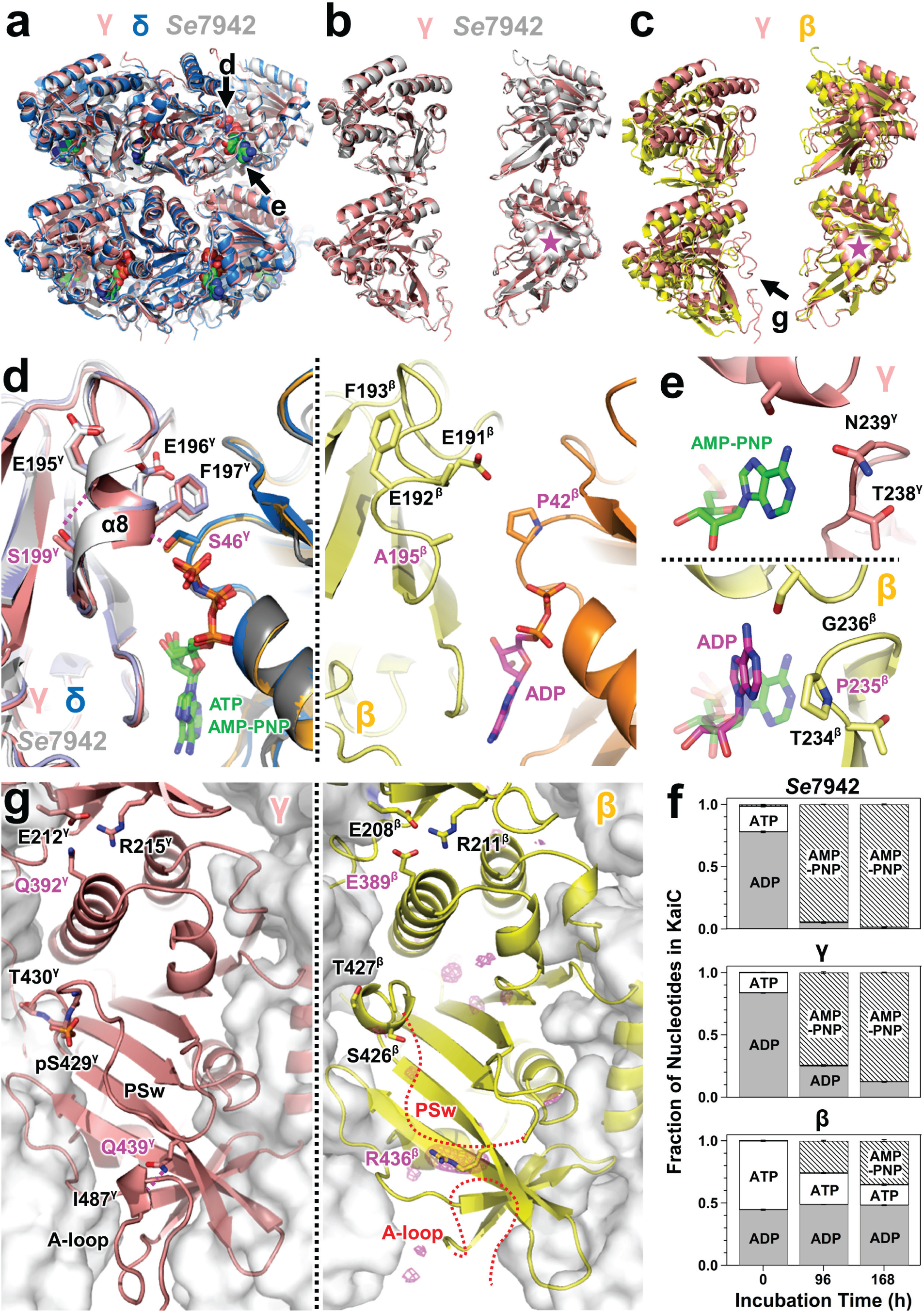
Crystal structures of ancestral KaiCs. (**a**) Similarity of the overall structures of KaiC^γ^ (pink), KaiC^δ^ (blue), and KaiC*^Se^* (gray, PDB ID: 7DXQ). Black thick arrows indicate the viewing directions for each panel. (**b**) Superimposition of KaiC^γ^ (pink) and KaiC*^Se^* (gray) using one of the CII domains (magenta star). The main-chain root-mean-squared deviation (RMSD) values of the fitted and other parts were 0.7 and 1.2 Å, respectively. For clarity, only the two most distantly located subunits within the hexamer are shown. (**c**) Superimposition of KaiC^γ^ (pink) and KaiC^β^ (yellow) using one of the CII domains (magenta star). Main-chain RSMD values of the fitted and other parts were 0.7 and 5.3 Å, respectively. (**d**) Zoomed-in views of the pre-hydrolysis CI–CI interfaces (left) for KaiC^γ^ (pink/orange), KaiC^δ^ (cyan/blue), and KaiC*^Se^* (white/gray), and the post-hydrolysis CI–CI interface (right) for KaiC^β^ (yellow/brown). Dotted magenta lines represent hydrogen bonds. (**e**) Zoomed-in views of the nucleotide enclosure loop for KaiC^γ^ (top) and KaiC^β^ (bottom). In bottom panel, KaiC^γ^ was superimposed on KaiC^β^ using one CI protomer, and then only the AMP–PNP molecule (green stick model) bound to KaiC^γ^ is shown for comparison with the ADP molecule (magenta stick model) of KaiC^β^. (**f**) Exchange with AMP–PNP was barely detected in KaiC^β^-bound ADP, in contrast to KaiC*^Se^*- and KaiC^γ^-bound ADP. Ancestral KaiCs and KaiC*^Se^* were incubated at pH 7.0 and 8.0, respectively, at 30°C in the presence of 1 mM AMP– PNP; the fractions of nucleotides bound by KaiCs were quantified. (**g**) CII domains of KaiC^γ^ (left) and KaiC^β^ (right) viewed from the inner-diameter side of the hexamers. The magenta meshes represent the *F*_obs_ – *F*_calc_ omit maps contoured at 3σ. Dotted red lines represent potentially flexible parts that could not be determined because of poor electron density.

Structural differences between KaiC^β^ and KaiC^γ^ were found in both the CI and CII domains. Three major differences were observed in the CI domain with the ATPase activity responsible for period-length determination ^29,30^. The first is the type of bound nucleotide. KaiC^γ^ and KaiC^δ^ were crystallized in the pre-hydrolysis state with a non-hydrolyzable ATP analog (AMP-PNP) bound to the CI domain, as described for KaiC*^Se^* ^29,39^ (left in Fig. 3d and Extended Data Fig. 6a); by contrast, KaiC^β^ was crystallized in the post-hydrolysis state with an adenosine diphosphate (ADP) bound to its CI active site even in the presence of 5 mM AMP–PNP (right in Fig. 3d and Extended Data Fig. 6b).

The second difference is the interface between two adjacent CI domains. In the pre-hydrolysis states of KaiC^γ^, KaiC^δ^, and KaiC*^Se^* (left in Fig. 3d), each CI–CI interface is stabilized by the hydrogen bond between the Ser residue side chain (S46^γ^, S44^δ^, and S48*^Se^*: Fig. 2a) in one CI domain and the carboxyl oxygen atom of the Glu residue (E196^γ^, E194^δ^, and E198*^Se^*) in the adjacent domain. However, in KaiC^β^, the corresponding Ser residue is replaced by Pro (P42^β^: Fig. 2a) and the packing of the CI–CI interface becomes loose because of the cleavage of the inter-domain hydrogen bond (right in Fig. 3d), as observed for the post-hydrolysis CI ring of KaiC*^Se^* ^27,29^. In addition, the replacement of the amino acid corresponding to S199 in KaiC^γ^ (S199^γ^) by A195 in KaiC^β^ (A195^β^) leads to further selective stabilization of the post-hydrolysis CI conformation of KaiC^β^. In the pre-hydrolysis state of KaiC^γ^ (left in Fig. 3d), the α_8_ helix is stabilized by the hydrogen bond between S199^γ^ and E195^γ^ (between S197^δ^ and E193^δ^ in KaiC^δ^, and between S201*^Se^* and E197*^Se^* in KaiC*^Se^*). In KaiC^β^, however, cleavage of this hydrogen bond by A195^β^ enables the partial unfolding of the α_8_ helix (right in Fig. 3d), as confirmed by the loose CI– CI interface of the post-hydrolysis CI of KaiC*^Se^*^27,29^. These observations support the preferential binding of ADP to the loose CI–CI interface of KaiC^β^.

The third difference is the insertion of a residue into the loop enclosing the nucleotide bound in the active site of the CI domain. Compared with KaiC*^Se^* and KaiC^γ^ (top in Fig. 3e), KaiC^β^ had a proline residue inserted at the 235^th^ position (P235^β^: Fig. 2a) and its structure showed that the residue was located in the turn of the nucleotide enclosure loop (bottom of Fig. 3e). The distorted structure of the ADP bound nearby suggests that, in KaiC^β^, P235^β^ pushes its adenine ring toward the center of the CI ring and inhibits the release of bound ADP. In rhythmic KaiCs such as KaiC^γ^, KaiC^δ^, and KaiC*^Se^*, no such inhibitory structures for releasing nucleotides exist. Consistent with this structural observation, biochemical experiments showed that ADP bound to KaiC^β^ was highly stabilized and, in contrast to ADP bound to KaiC*^Se^* and KaiC^γ^, was not easily exchanged by AMP–PNP (Fig. 3f).

These observations suggest that the loose CI–CI interface results in the preferential binding of ADP (right in Fig. 3d) and that the elongation of the nucleotide enclosure loop (bottom in Fig. 3e) further prevents the exchange of generated or bound ADP (Fig. 3f). These factors might have caused the consistent occupation of the CI site by ADP in KaiC^β^, resulting in a reduction of the ATPase activity (1.8 ± 0.2 ATP d^-1^ at 30°C, Fig. 2b) and the loss of the damped oscillatory relaxation in KaiC^β^ (Fig. 2c). KaiC might have acquired its oscillatory ability by evolving its CI structure to enable nucleotides bound to the CI domain to be exchanged on a reasonable circadian time scale.

The structural differences between KaiC^β^ and KaiC^γ^ were not only observed in the CI–CI interface but also in the CII domain and the CI–CII interface. In rhythmic KaiC^γ^, a region upstream of the C-terminal tail to which KaiA^γ^ binds ^40^ was clearly resolved as a loop structure (A-loop) with a stabilizing hydrogen bond between Q439^γ^ and I487^γ^ (left in Fig. 3g), as observed between Q441*^Se^* and I489*^Se^* in KaiC*^Se^* ^39,41^. In KaiC^β^, however, the A-loop was destabilized by R436^β^, which breaks this hydrogen bond (right in Fig. 3g). The electron density of the phosphor-switch, which changes conformation upon phosphoryl modification of the dual phosphorylation site (S431*^Se^* and T432*^Se^*) in rhythmic KaiC*^Se^* ^39^, was clearly identified in KaiC^γ^ but not in KaiC^β^ (right in Fig. 3g). In addition, the 389^th^ residue located in the CI–CII interface is Glu in KaiC^β^ (E389^β^), whereas the corresponding residue is Gln in the rhythmic KaiC^γ^ (Q392^γ^) and KaiC*^Se^* (Q394*^Se^*). In KaiC*^Se^*, Q394*^Se^* is critical for establishing the rhythmic coupling between the CI and CII domains by switching the interaction with E214*^Se^*and/or R217*^Se^* (Fig. 2a and left in Fig. 3g), as evidenced by the arrhythmicity of the Q394E variant of KaiC*^Se^* ^39^.

## Discussion

### A possible scenario for the evolution of the Kai protein oscillator

The 16S rRNA–based phylogenic tree (Extended Data Fig. 2) shows the presence of a node (γ point) diverging into freshwater and marine cyanobacteria, which can be interpreted as corresponding to the MRCA of cyanobacteria. KaiC^γ^ is the topologically identical node that branches into the extant KaiCs of freshwater cyanobacteria (group I) and marine cyanobacteria (group II) (Fig. 1a and Extended Data Fig. 1), and the same applies to KaiB^γ^ (Extended Data Fig. 4) and KaiA^γ^ (Extended Data Fig. 5). These observations clearly indicate that the A^γ^B^γ^C^γ^ set functioned as the self-sustained circadian oscillator in the MRCA of cyanobacteria capable of oxygenic photosynthesis.

There are many estimates as to when the cyanobacterial MRCA first appeared ^2^, which range from 3.5 Ga to 1.5 Ga ago ^1,3-6^. Our estimate obtained using 16S rRNA sequences dated an ancestral bacterium at an α/β point at 3.1 (3.5–2.8: upper–lower confidence intervals) Ga ago and the MRCA of cyanobacteria at the γ point at 2.2 (2.8–1.7) Ga ago (Fig. 4 and Extended Data Table 3). An ancestral form of the double-domain KaiA, which is evolutionarily related to KaiA^γ^, might have appeared approximately 1.0 Ga ago ^32^; however, its appearance was subsequently updated to an earlier period, as late as 1.5 Ga ago ^36^.

**Fig. 4.**
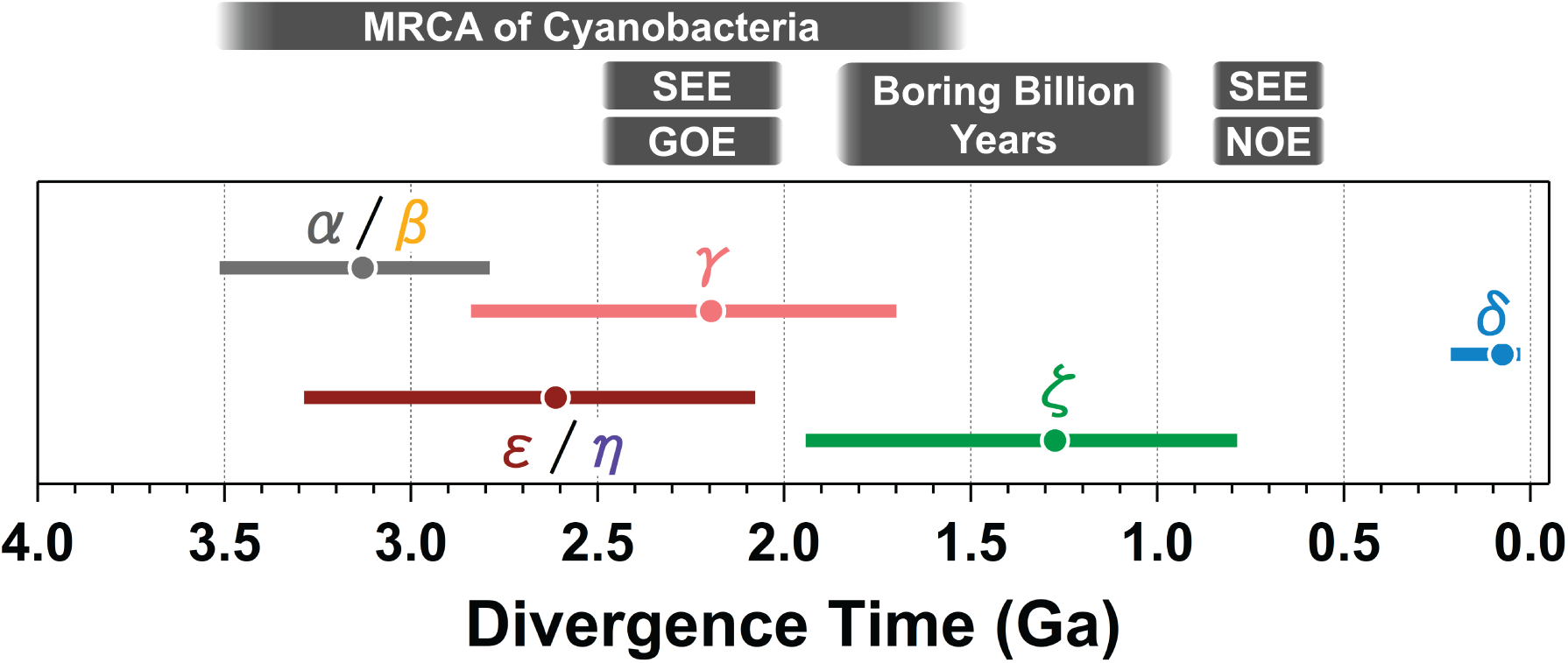
Evolutionary origins and diversification of self-sustained Kai protein circadian oscillators. Divergence times were estimated on the basis of TimeTree analysis using 16S rRNA sequences. Filled circles and horizontal bars indicate estimated divergence times (1 Ga = 1 billion years) and confidence intervals, respectively (Extended Data Table 3). The approximate times of the emergence of the most recent common ancestor (MRCA) of cyanobacteria ^1-6^, Great Oxidation Event (GOE) ^7^, first and second Snowball Earth events (SEEs) ^8-10^, the post-GOE period known as the "boring billion years" of ever-stable global conditions ^58^, and the Neoproterozoic Oxygenation Event (NOE) ^11,12^ are schematically shown for reference.

Under our divergence time estimate, the MRCA of cyanobacteria would have emerged around the GOE and first SEE period (Fig. 4). This period is characterized by the enhancement of oxygenic photosynthesis due to changes in the properties of ancient cyanobacteria ^1-3^ or other ancient bacteria ^4-6^, which destroyed the methane greenhouse and triggered global glaciation ^42^. The resonance between the endogenous clock and environmental light–dark cycles is the basis for more efficient growth and survival of organisms, including cyanobacteria ^43^. Thus, it is reasonable to speculate that the acquisition of the autonomous γ set allowed the MRCA of cyanobacteria to expect the timing of the next sunrise and prepare in advance to initiate oxygenic photosynthesis accordingly. The Earth’s rotation period 2 Ga ago has been suggested to be 2–4 h shorter than that at present ^14^; however, depending on the pH of the growing environment, the γ set in the cyanobacterial MRCA might have been sufficiently functional as a circadian oscillator (Fig. 1k).

Regardless of the accuracy of the divergence time estimation, another ancestral bacterium corresponding to the ε/η point appeared at approximately the same time as the MRCA of cyanobacteria (Fig. 4). The ε/η point is assumed to possess not–self-sustained (Fig. 1o) and temperature-dependent (Fig. 2b) KaiC^η^, an ancestral form of the hourglass-like KaiC*^Rs^* ^34^ and not– self-sustained KaiC*^Rp^* ^35^. The self-sustained protein-based circadian oscillator established in the γ set was subsequently inherited by KaiC^ζ^ and KaiC*^Se^* and, consequently, by approximately 70% of group I cyanobacteria in freshwater (Fig. 1a and Extended Data Fig. 1). The inheritance from KaiC^γ^ applied not only to KaiC^ζ^ but also to KaiC^δ^, and the autonomous oscillatory property was eventually passed on to marine cyanobacteria located downstream of KaiC^δ^. However, our results indicate that it was not inherited by the *Prochlorococcus* group located downstream of KaiC^δ^ or that the inherited traits have changed by today. Given that the molecular evolution of oxygenic photosynthesis pre-dates the cyanobacterial MRCA by >1 Ga ^2,4^, the unique evolution of the ancestral Kai proteins during the emergence of the cyanobacterial MRCA provided the autotrophic basis for the time-optimal acquisition and consumption of energy, later leading to their massive proliferation across various habitats on Earth.

## Methods

### Construction of protein-based phylogenic trees

The amino acid sequences of KaiC homologs were obtained from a dataset described previously ^33^. From a total of 1,194 entries, duplicate entries that had identical sequences but were from different species/strains were removed, resulting in 151 entries of 480–570 amino acid residues with >30% sequence identity to KaiC*^Se^*. These were extracted to form a dataset for analyzing KaiC homologs with double-domain structures (Extended Data Fig. 1).

The MEGA X software ^44^ was used for phylogenic analyses. The 151 sequences were arranged using pairwise/multiple alignments with a gap-opening penalty of 7.5. A preliminary phylogenetic tree was constructed using the maximum likelihood method, and 100 bootstrap replications were generated. To confirm the stability of the resultant phylogenetic pattern, different alignment results were analyzed using gap-opening penalties of 2.5, 5.0, 10.0, 15.0, and 20.0. In the preliminary phylogenetic trees, double-domain homologs of KaiC were classified into four groups as follows: cyanobacteria in freshwater (group I: 58 entries), cyanobacteria in seawater (group II: 26 entries), other bacteria (group III: 23 entries), other bacteria including archaea (group IV: 42 entries), and outgroup (2 entries). The final phylogenetic tree of double-domain KaiC homologs (Fig. 1a) was obtained using a gap-opening penalty of 7.5 through 500 bootstrap replications. See Supplementary Information and Extended Data Fig. 1 for a detailed description of the entries in each group.

Phylogenetic analyses of KaiA and KaiB homologs were performed as described for KaiC using a dataset from a previous study ^33^. For KaiB, we extracted 193 entries of 80–120 amino acid residues with >30% sequence identity to KaiB*^Se^* (Extended Data Fig. 4). For KaiA, extraction was performed with 50–350 amino acid residues with >30% sequence identity to KaiA*^Se^*, resulting in 151 entries of double-domain homologs of KaiA (Extended Data Fig. 5). The species used to construct the phylogenetic trees for KaiA and KaiB were not necessarily identical to those used for the KaiC tree. The final phylogenetic trees for KaiB (Extended Data Fig. 4) and KaiA (Extended Data Fig. 5) were obtained using a gap-opening penalty of 7.5 through 500 bootstrap replications. See Supplementary Information for detailed descriptions of the entries in each group.

### Construction of 16S rRNA–based phylogenic trees

The 16S rRNA sequences of extant species with double-domain KaiC homologs were obtained from KEGG ^45^. The 16S rRNA–based tree shown in Extended Data Fig. 2 was constructed using MEGA X software with 108 entries of the 151 KaiC homologs used for construction of the protein-based tree (Fig. 1a).

### Restoration of amino acid sequences of ancestral KaiCs

The amino acid sequences corresponding to seven nodes on the phylogenic tree of KaiC (Fig. 1a) were inferred on the basis of the maximum likelihood estimation using the amino acid substitution model of LG with Freqs. (+F) and gamma-distributed substitution rates. The oldest ancestor (KaiC^α^) was detected directly below a node branching into groups I–IV, *Pyrococcus horikoshii* OT3 KaiC (KaiC*^Hori^*), and *Pyrodictium delaneyi* KaiC (KaiC*^Dela^*).

There were two branches emerging from KaiC^α^: one ended in groups I–III, and the other ended in group IV. A node connecting to groups I–III (KaiC^β^) showed high bootstrap values (98%– 99%) irrespective of the gap penalties (7.5–15). The KaiC^β^ node branched into groups I–II and group III. A node branching into groups I and II (KaiC^γ^) also emerged with high bootstrap values (99%) irrespective of the gap penalties used. Although the common ancestor of group II (KaiC^δ^) showed a high bootstrap value (31%–97%), those of group I (KaiC^ζ^) and group III (KaiC^ε^) showed relatively lower values (20%–51% and 53%–63%, respectively). The common ancestor of group IV (KaiC^η^) was detected with the bootstrap vale (98%) as high as those of KaiC^β^ and KaiC^γ^.

### Restoration of amino acid sequences of ancestral KaiAs and KaiBs

The amino acid sequences corresponding to the nodes on the phylogenic trees of KaiA and KaiB were estimated as described for ancestral KaiCs. In the KaiB tree constructed with 193 entries (Extended Data Fig. 4), we assigned one node for each, KaiB^γ^ and KaiB^δ^, on the basis of similarities in the taxonomic patterns of the corresponding entries (see coloring by growth habitat in Extended Data Fig. 1 and Extended Data Fig. 4). The KaiB^δ^ node was located downstream of KaiB^γ^. Because a node upstream of KaiB^γ^ was a common ancestor for all entries, it was termed KaiB^α/β^. A node that did not contain downstream archaeal entries was termed KaiB^ε^.

Only three later branching points (γ, δ, and ζ) could be assigned in the double-domain KaiA tree consisting of 151 entries (Extended Data Fig. 5), which is consistent with the later appearance of ancestral KaiA than ancestral KaiB ^36^. KaiA^γ^ was reasonably allocated to a node corresponding to a common ancestor of the double-domain KaiA from both freshwater cyanobacteria and seawater cyanobacteria. Similarly, KaiA^ζ^ and KaiA^δ^ were assigned to nodes corresponding to ancestors of cyanobacteria in freshwater and seawater, respectively.

### Divergence time estimation

The divergence times of the α/β, γ, δ, ε/η and ζ points in the 16S rRNA–based tree (Extended Data Fig. 2) were estimated using MEGA X software ^44^. According to the TimeTree ^46^ estimate of the divergence time between *Synechococcus* and *Flavobacterium*, a calibration point was established at the α/β point as normal distribution with a median at 3.134 Ga and a deviation of 0.5 Ga.

### Cloning of extant and ancestral *kaiA*, *kaiB*, and *kaiC*

The DNA sequences of extant and ancestral Kai-proteins were synthesized by Eurofins genomics with codon optimization for expression in *E. coli*. Unless otherwise stated, the synthesized genes were cloned into the pGEX6P-1 vector (Cytiva) with *BamHI* and *NotI* cleaving sites to overexpress GST-tagged Kai proteins. For ancestral KaiC^α^, KaiC^β^, KaiC^ε^, and KaiC^η^, the synthesized genes were cloned into the pET-3a vector (Novagen) with *NdeI* and *BamHI* cleaving sites using nucleotides encoding hexagonal histidine (His) at the 3′ end. For extant KaiCs from *Flavobacterium johnsoniae* UW101 and *Rhodopseudomonas palustris* TIE-1 strains, genes were cloned into the pASK-IBA5plus vector (IBA GmbH) with the *BsaI* cleaving site to overexpress Strep-tagged KaiCs because of the poor yields associated with the GST-tagged form.

### Expression and purification of extant and ancestral Kai-proteins

Extant and ancestral KaiCs were expressed and purified as described previously ^47^ according to the type of fusion tags used. Irrespective of the extant or ancestral types, KaiA and KaiB were purified following the methods used for KaiA*^Se^* and KaiB*^Se^*, respectively ^48^.

### *In vitro* rhythm assay

For the *in vitro* rhythm assay of the extant species, KaiA, KaiB, and KaiC were mixed in Tris-buffer (20 mM Tris, 150 mM NaCl, 5 mM MgCl_2_, and 0.5 mM EDTA, pH 8.0) containing 1 mM ATP, and the time-evolution of the phosphorylation of KaiC was monitored. The potential oscillation of the respective KaiCs was assessed in the presence of various concentrations of KaiA (0.01, 0.02, 0.04, and 0.08 mg/mL) and a fixed concentration of KaiB (0.04 mg/mL) according to previous observations that the amount of KaiA in the system markedly affects the amplitude or stability of the rhythm ^31,49,50^. KaiCs that showed any slight sign of oscillation under either condition exhibited a stable rhythm in the presence of moderate concentrations of KaiA (0.02– 0.04 mg/mL) (Extended Data Fig. 3). For the assay of KaiC*^Cy2^* from *Cyanothece sp.* ATCC 51142 (ID: 085), a KaiA homolog (protein ID: WP_009543244.1, ∼17% identity to KaiA*^Se^*) that is not listed in the present KaiA tree but has recently been reported as promising was used as the species-specific KaiA ^51^.

For the *in vitro* rhythm assay of the ancestral forms, KaiA, KaiB, and KaiC were mixed Tris-buffer (20 mM Tris, 150 mM NaCl, 5 mM MgCl_2_, and 0.5 mM EDTA, pH 7.0) containing 1 mM ATP at the following final concentrations (mg/mL);

A^γ^ : B^α/β^ : C^α^ = 0.04 : 0.04 : 0.2;
A*^Se^* : B*^Se^* : C^α^ = 0.04 : 0.04 : 0.2;
A^γ^ : B^α/β^ : C^β^ = 0.04 : 0.04 : 0.2;
A*^Se^* : B*^Se^* : C^β^ = 0.04 : 0.04: 0.2;
A^γ^ : B^γ^ : C^γ^ = 0.04 : 0.04 : 0.2;
A*^Se^* : B*^Se^* : C^γ^ = 0.04 : 0.04 : 0.2;
A^δ^ : B^δ^ : C^δ^ = 0.04 : 0.04 : 0.2;
A*^Se^* : B*^Se^* : C^δ^ = 0.04 : 0.04 : 0.2;
A^γ^ : B^ε^ : C^ε^ = 0.04 : 0.04 : 0.2;
A*^Se^* : B*^Se^* : C^ε^ = 0.04 : 0.04 : 0.2;
A^ζ^ : B^γ^ : C^ζ^ = 0.05 (30°C) or 0.06 (40°C) : 0.02 : 0.2;
A*^Se^* : B*^Se^* : C^ζ^ = 0.04 : 0.04 : 0.2;
A^γ^ : B^η^ : C^η^ = 0.04 : 0.04 : 0.2; and
A*^Se^* : B*^Se^* : C^η^ = 0.04 : 0.04 : 0.2.

### ATPase measurements

ATPase measurements were performed using an ultrahigh-performance liquid chromatography (UHPLC) system (Chromaster, Hitachi), as reported previously ^47^. Ancestral KaiCs and KaiC*^Se^* were incubated in a Tris-buffer containing 1 mM ATP at pH 7.0 and pH 8.0, respectively. The ATPase activity of KaiC was defined as the number of hydrolyzed ATP molecules into ADP molecules per KaiC monomer per unit time. Statistical differences in the ATPase activities between 30°C and 40°C were analyzed using ANOVA and the Bonferroni post hoc *t* test.

### Pre–steady-state ATPase assay

For the pre–steady-state ATPase assay ^29^, an apo-KaiC monomer was first prepared as described previously ^52^ using a phospho-buffer (50 mM NaH_2_PO_4_, 150 mM NaCl, 50 mM L-arginine, 50 mM L-glutamic acid, 5 mM MgCl_2_, and 1 mM DTT, pH 7.8). Reassembly of KaiC hexamers was induced by mixing the apo-KaiC monomer with the phospho-buffer containing ATP, and the resultant mixture containing 1 mM ATP was immediately subjected to UHPLC analysis for ATPase measurements.

### *In vivo* rhythm assays

*Synechococcus elongatus* PCC 7942 was used as the background strain for this assay. The *in vivo* bioluminescence assay and analysis were performed as reported previously ^16^. Cyanobacterial cells carrying the *kaiBC*-reporter cassette were cultured at 30°C for 3 days on BG-11 solid medium under constant light (LL) at 100 μmol m^−2^ s^−1^. After one or three 17-h light / 17-h dark treatments, the cells were cultured at 30°C under LL (45 μmol m^−2^ s^−1^) at an LED daylight lamp, and bioluminescence profiles were monitored using a photomultiplier tube detector.

### X-ray structure analyses of KaiC^β^, KaiC^γ^, and KaiC^δ^

Crystals of KaiC^β^, KaiC^γ^, and KaiC^δ^ were obtained using the sitting-drop vapor diffusion method with 5 mM AMP-PNP. KaiC^β^ (3 mg/mL) was mixed with a solution of 18 mM acetic acid and 65 mM sodium malonate (pH 5.7). The cryo-protectant treatment was performed using 30% (w/v) PEG4K. KaiC^γ^ (3 mg/mL) was crystallized in a solution of 0.94 M sodium formate and 1.2 M sodium malonate (pH 8.5), which are both cryo-protectants. KaiC^δ^ (3 mg/mL) was added to 0.1 M Tris-HCl (pH 7.0), 1 M KCl, and 1.14 M sodium malonate.

KaiC^β^, KaiC^γ^, and KaiC^δ^ crystals were frozen using liquid nitrogen and placed under the cryo-stream of the SPring-8 BL44XU beamline (Harima, Japan). The crystals were exposed to X-ray irradiation at a wavelength of 0.9 Å, and diffraction images were collected by using an EIGER X 16M detector (Dectris) and the semi-automatic data reduction software, KAMO ^53^. Diffraction intensity data were processed using XDS ^54^. The initial models for KaiC^β^, KaiC^γ^, and KaiC^δ^ were obtained using the molecular replacement method with MOLREP ^55^ and the coordinates of the KaiC*^Se^*hexamer (PDB ID: 2GBL) as a template. Refinement was achieved using REFMAC5 ^56^. The model was constructed by using COOT ^57^. PyMOL (Schrödinger) was used for the graphic presentation of the refined models.

The KaiC^β^, KaiC^γ^, and KaiC^δ^ crystals had lattices of *P*2_1_ (*a* = 93 Å, *b* = 386 Å, *c* = 108 Å, β = 113°; two hexamers in an asymmetric unit), *P*1 (*a* = 91 Å, *b* = 110 Å, *c* = 167 Å, α = 78°, β = 87°, γ = 82°; single hexamer in an asymmetric unit), and *P*2_1_ (*a* = 92 Å, *b* = 175 Å, *c* = 95 Å, β = 97°; single hexamer in an asymmetric unit), respectively.

## Data Availability

All data needed to evaluate the conclusions in the paper are provided in the paper and/or the Supplementary Information. The atomic coordinates and structure factors have been deposited in the Protein Data Bank with accession codes 8ZN5 (KaiC^β^), 8ZN6 (KaiC^γ^), and 8ZN7 (KaiC^δ^).

## Acknowledgements

Diffraction data were collected at beamline BL44XU at the SPring-8 facility under the proposals 2021A6700, 2021B6602, 2023A6500, 2023B6842, and 2024A6930. This research was partly supported by the Platform Project for Supporting Drug Discovery and Life Science Research (BINDS) from AMED under grant number JP21am0101072 (support number 0583). This study was supported by Grants-in-Aid for Scientific Research (22H04984 to S.A., A.M., and Y.F., 17H06165 to S.A. and Y.F., 24H02301 to S.A., 22K15051 to Y.F., 19K16061 to Y.F., and 22K06175 to A.M.), Takeda Science Foundation (to S.A.), and Toyoaki Scholarship Foundation (to S.A.).

## Author Contributions

A.M., Y.F., and S.A. conceived the project. S.A., A.M., and Y.F. supervised the project. A.M. and Y.F. performed most of the biochemical experiments and analysis. A.M., Y.F., Y.O., and K.H. performed rhythm and ATPase assays. Y.O. performed the nucleotide-exchange experiments. Y.F. collected the diffraction data and analyzed it with input from E.Y. K.I.-M. performed *in vivo* experiments. S.A., A.M., and Y.F. drafted the manuscript. All the authors read and commented on the manuscript.

## Extended Data Figures and Tables

**Extended Data Fig. 1.**
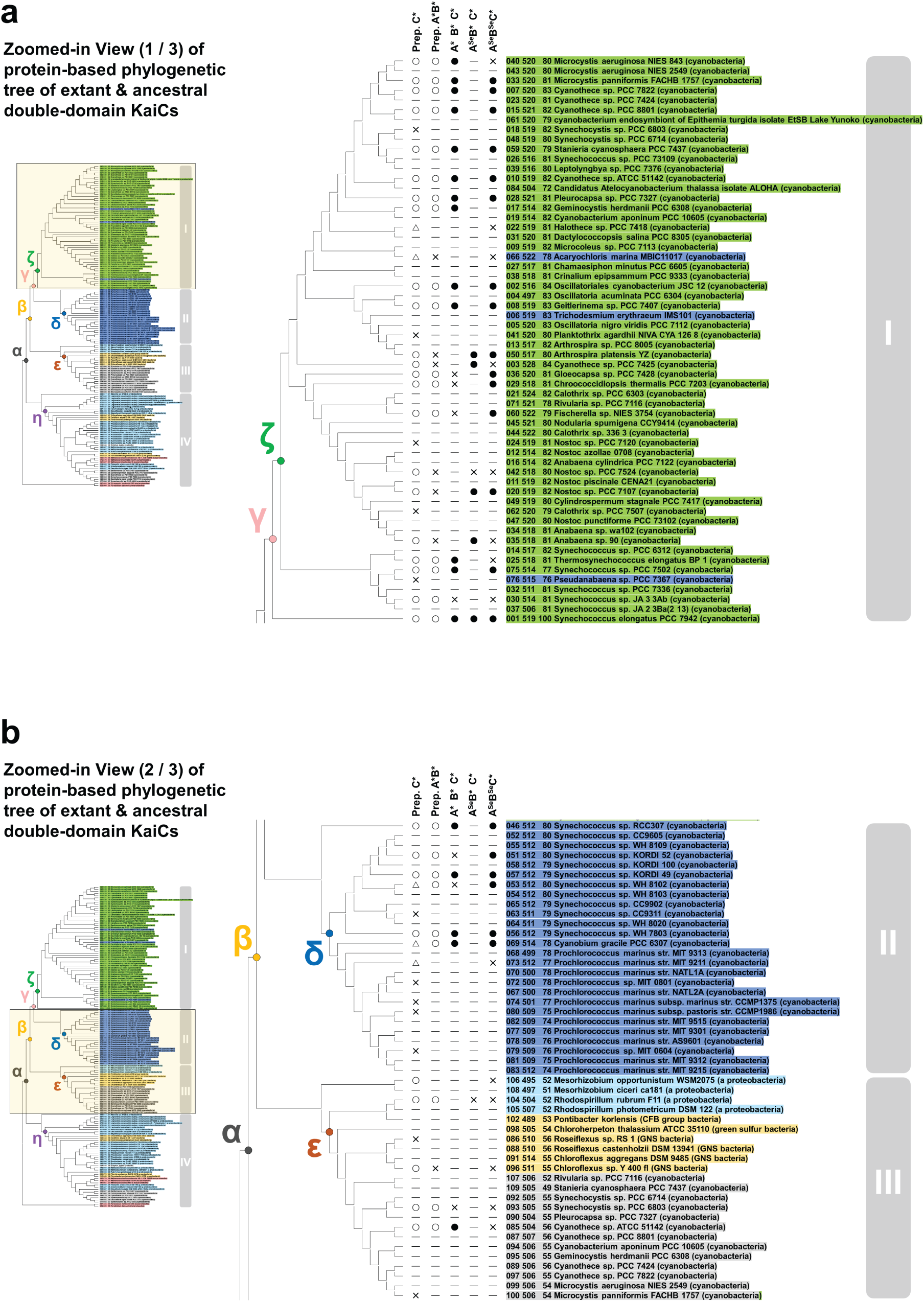
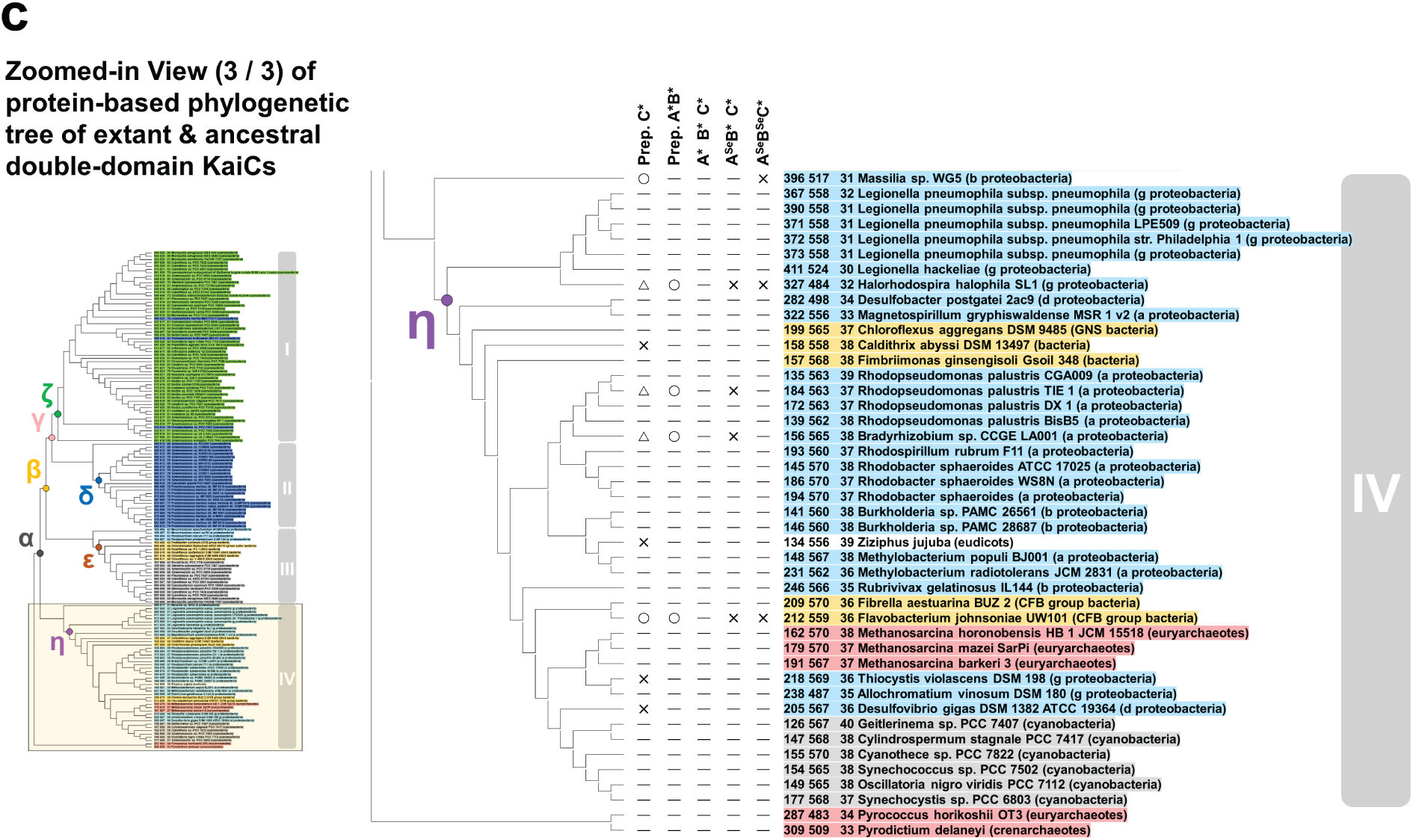
Zoomed-in views of the protein-based phylogenetic tree of extant and ancestral double-domain KaiCs. In each panel (**a**, **b**, and **c**), the enlarged image in the center corresponds to the area surrounded by a yellow square in the overall phylogenetic tree shown on the left. Cyanobacteria in freshwater, marine cyanobacteria, proteobacteria, bacteria, and euryarchaeota/crenarchaeota are colored green, blue, cyan, yellow, and red, respectively. Group III and IV KaiCs from freshwater cyanobacteria (gray entries) are originated from secondary copies of group I *kaiC* gene or lateral gene transfer, and thus can be excluded when identifying groups I–IV and nodes based on species and growing habitant. The three numbers preceding the name of each entry indicate the ID number, the total number of amino acids, and the sequence identity with KaiC*^Se^*, respectively. Refer to the caption of Fig. 1a for the notation method in the five items (Prep. C*, Prep. A*B*, A*B*C*, A*^Se^*B*C*, and A*^Se^*B*^Se^*C*).

**Extended Data Fig. 2.**
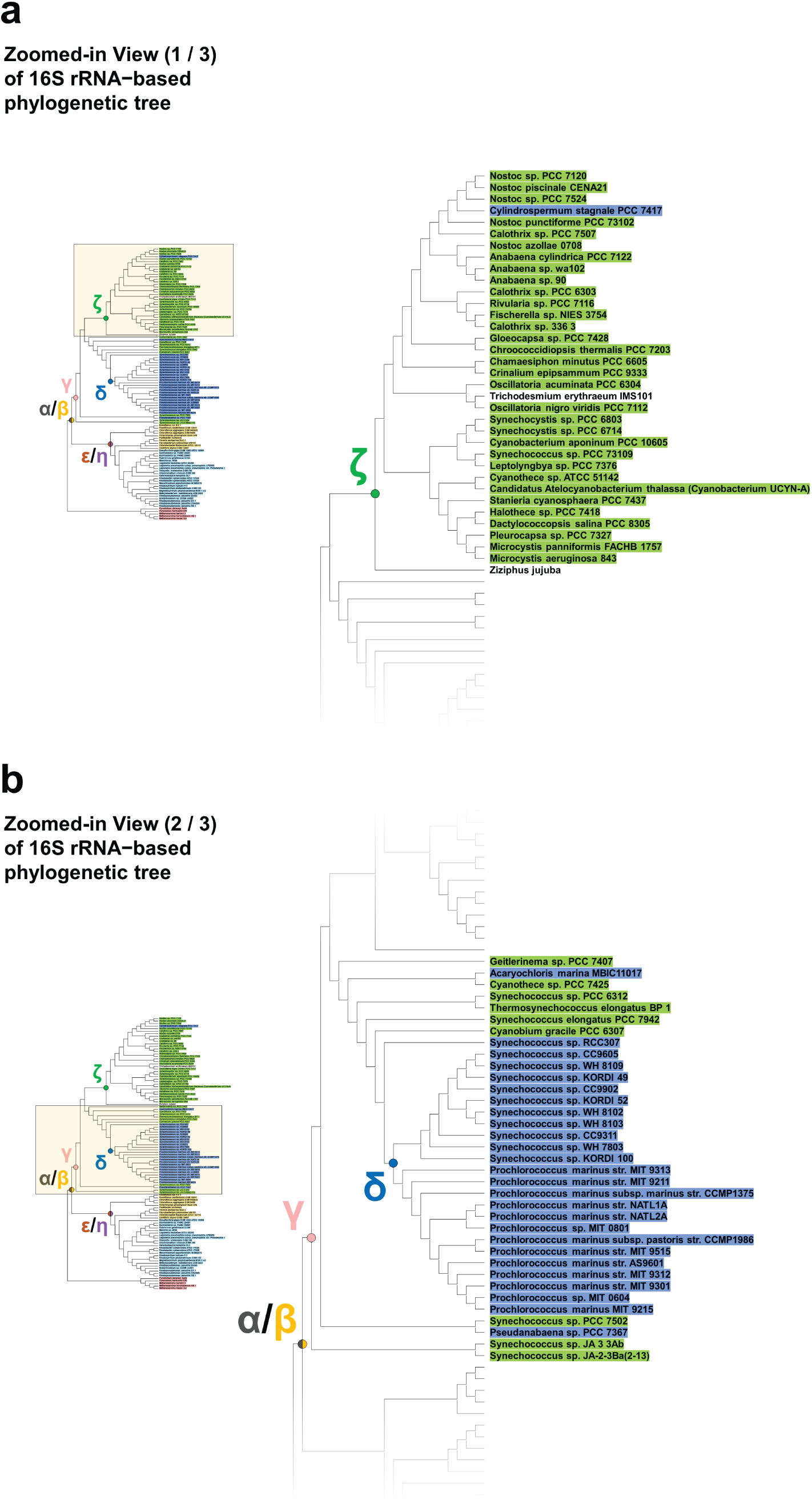
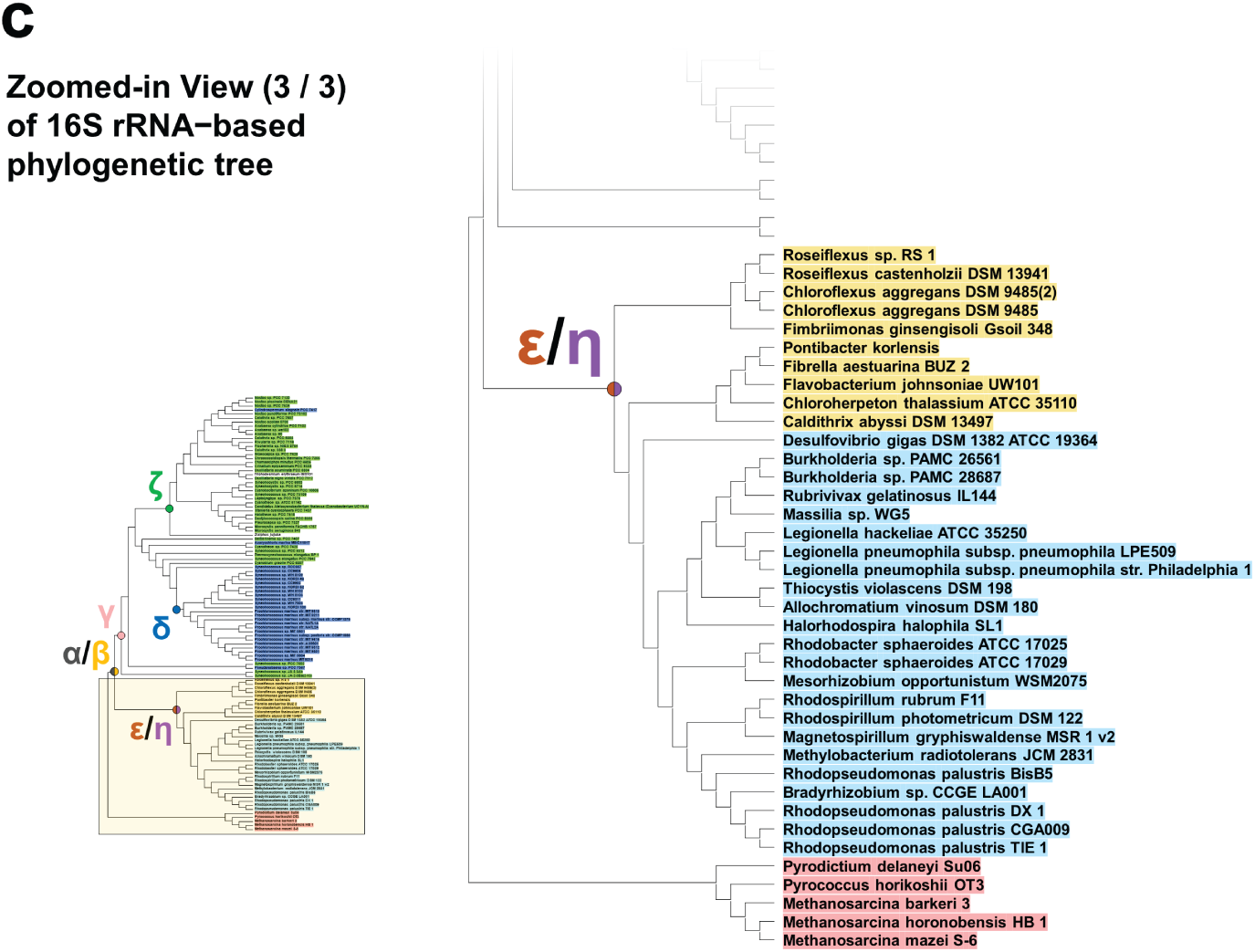
16S rRNA–based phylogenic tree. In each panel (**a**, **b**, and **c**), the enlarged image in the center corresponds to the area surrounded by a yellow square in the overall phylogenetic tree shown on the left. The 16S rRNA sequences were obtained from 108 entries of the 151 KaiC homologs used for protein-based tree construction (Fig. 1a and Extended Data Fig. 1). Cyanobacteria in freshwater, marine cyanobacteria, proteobacteria, bacteria, and euryarchaeota/crenarchaeota are colored green, blue, cyan, yellow, and red, respectively.

**Extended Data Fig. 3.**
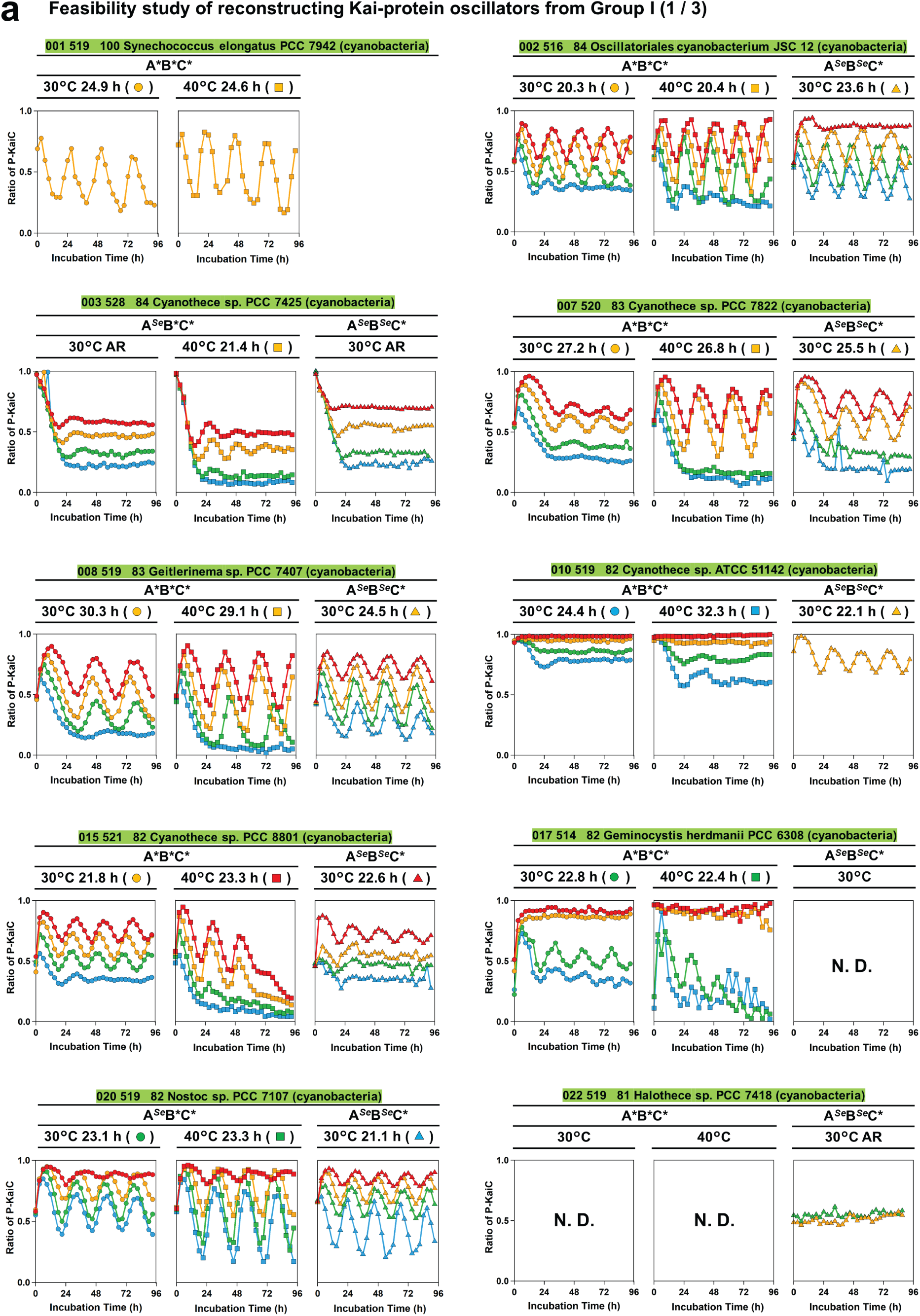
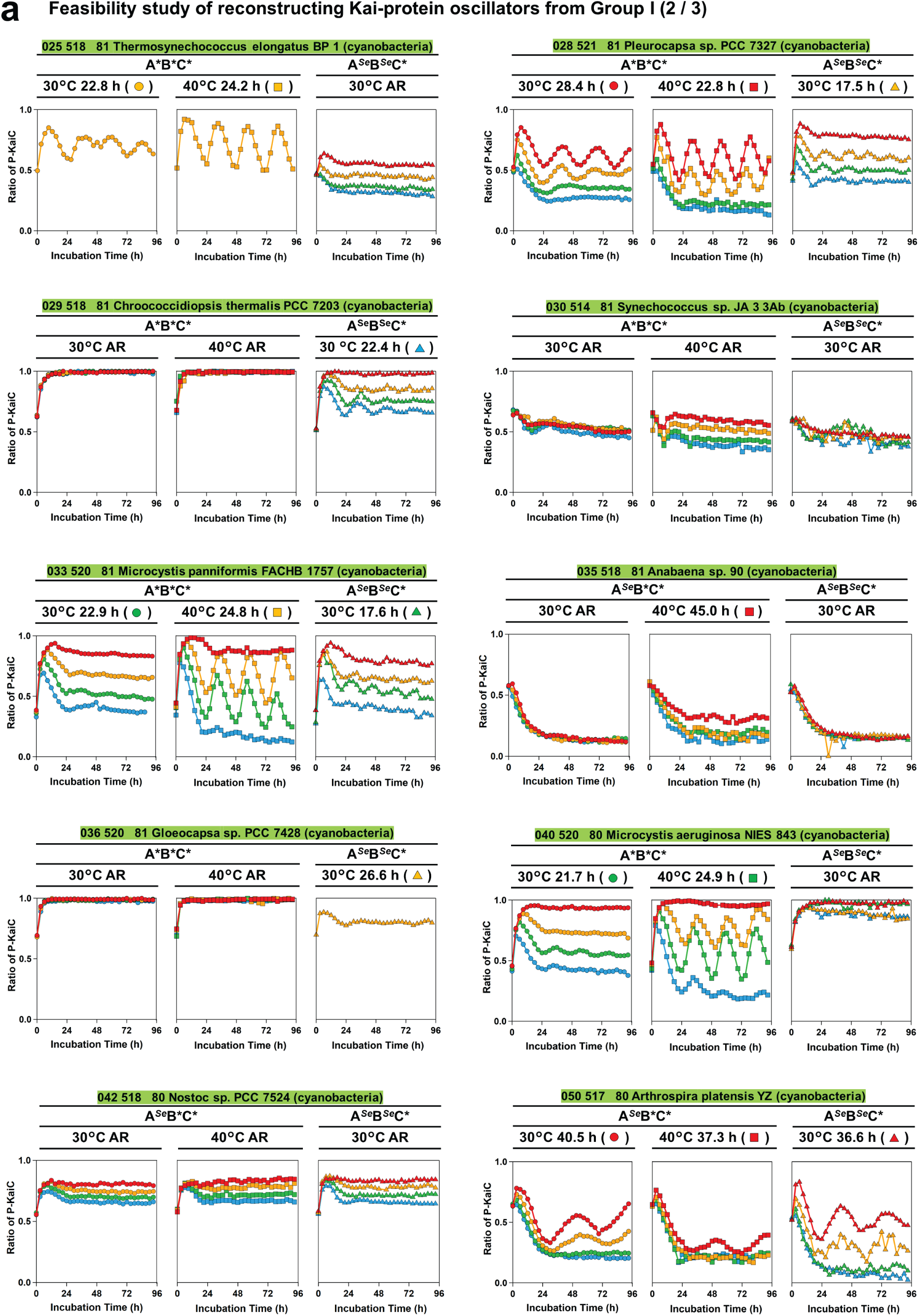
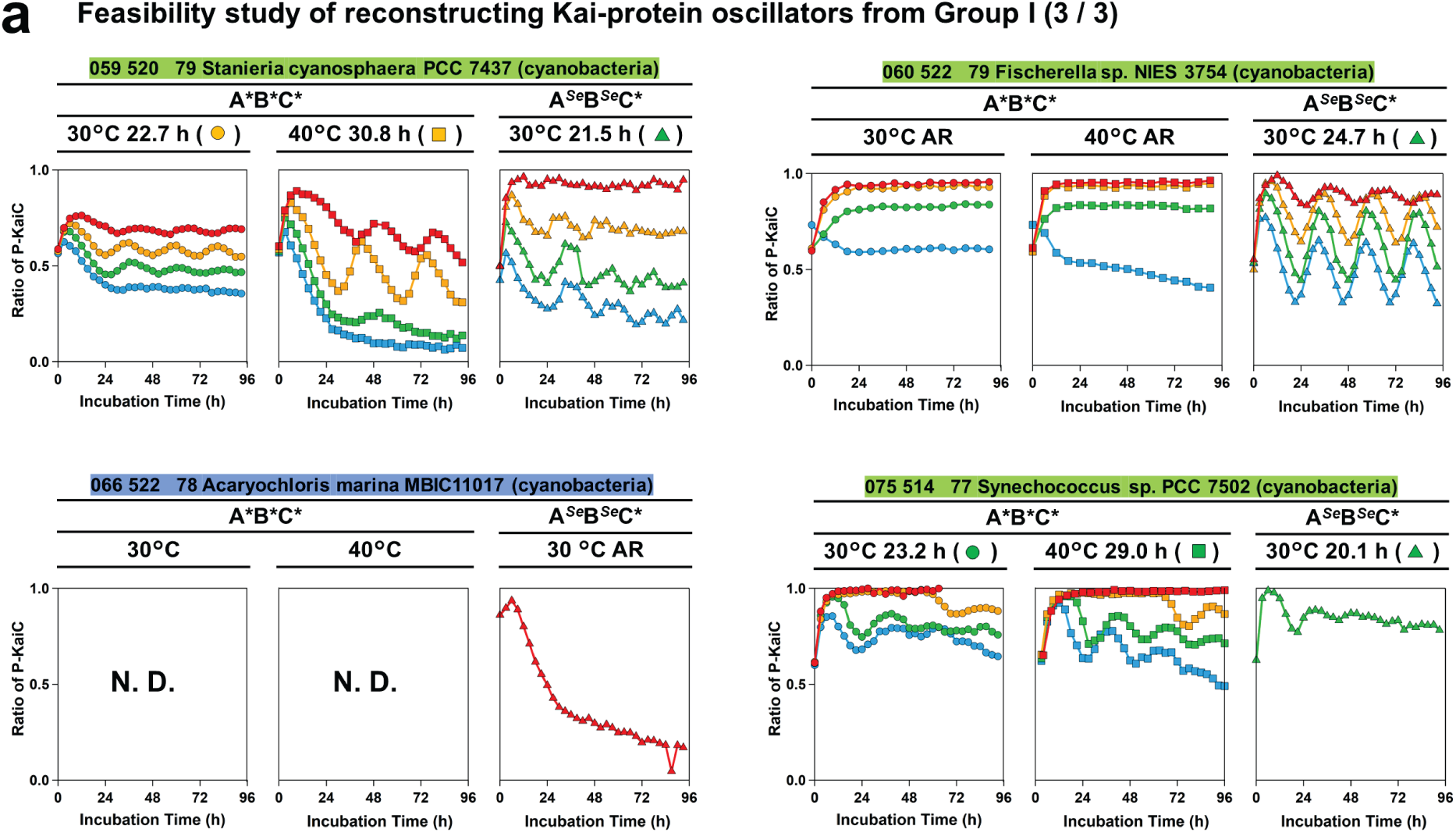
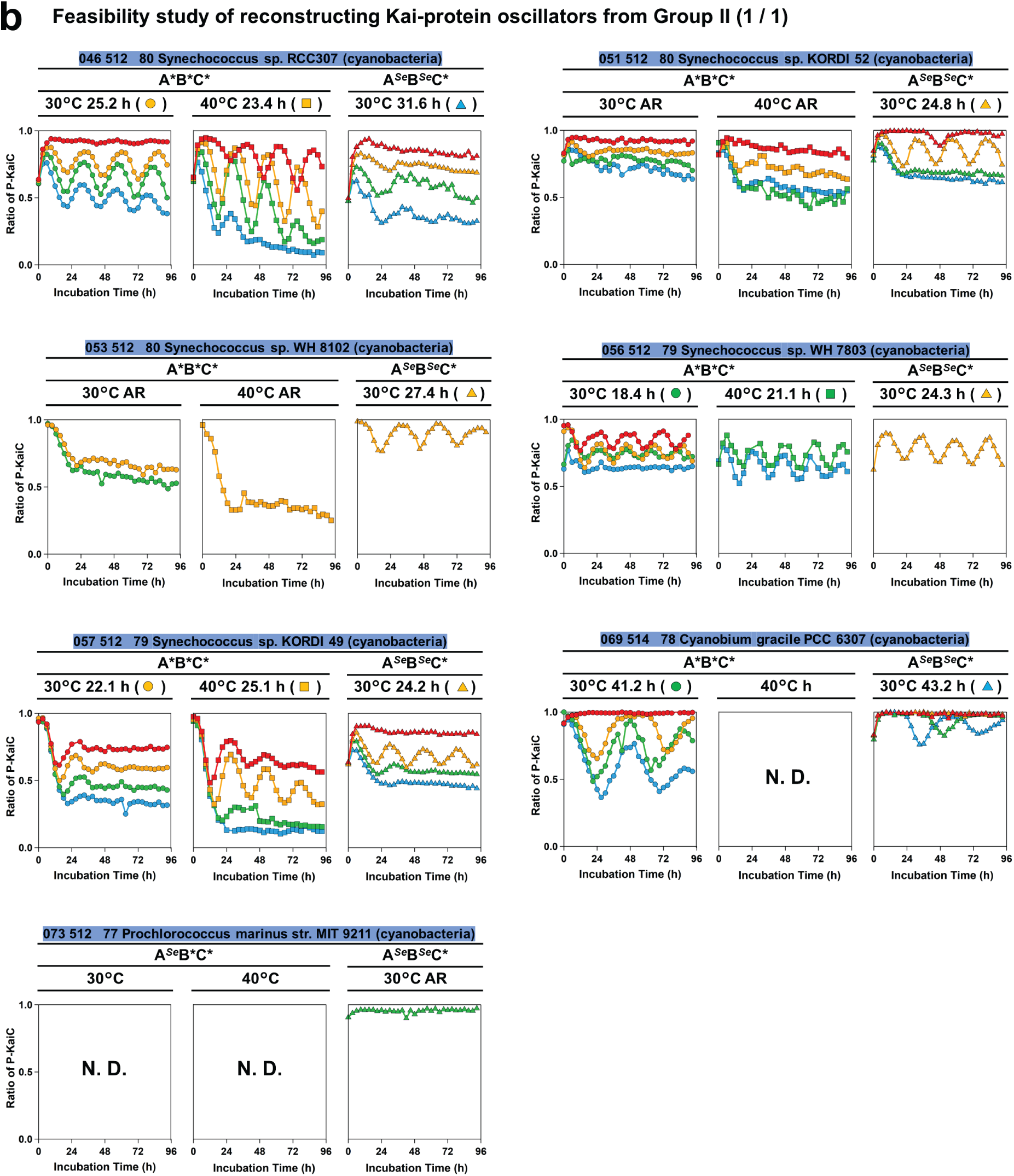
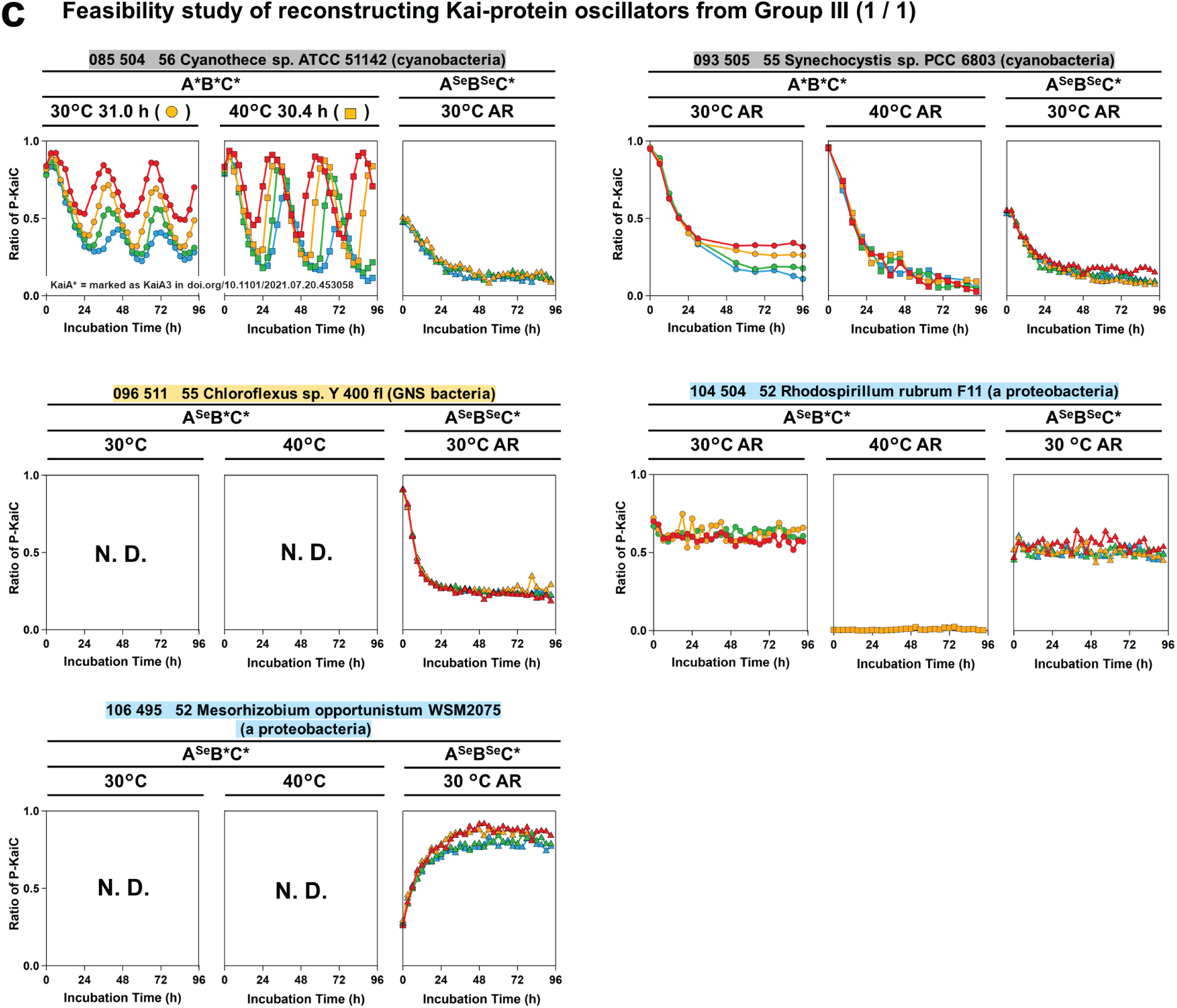
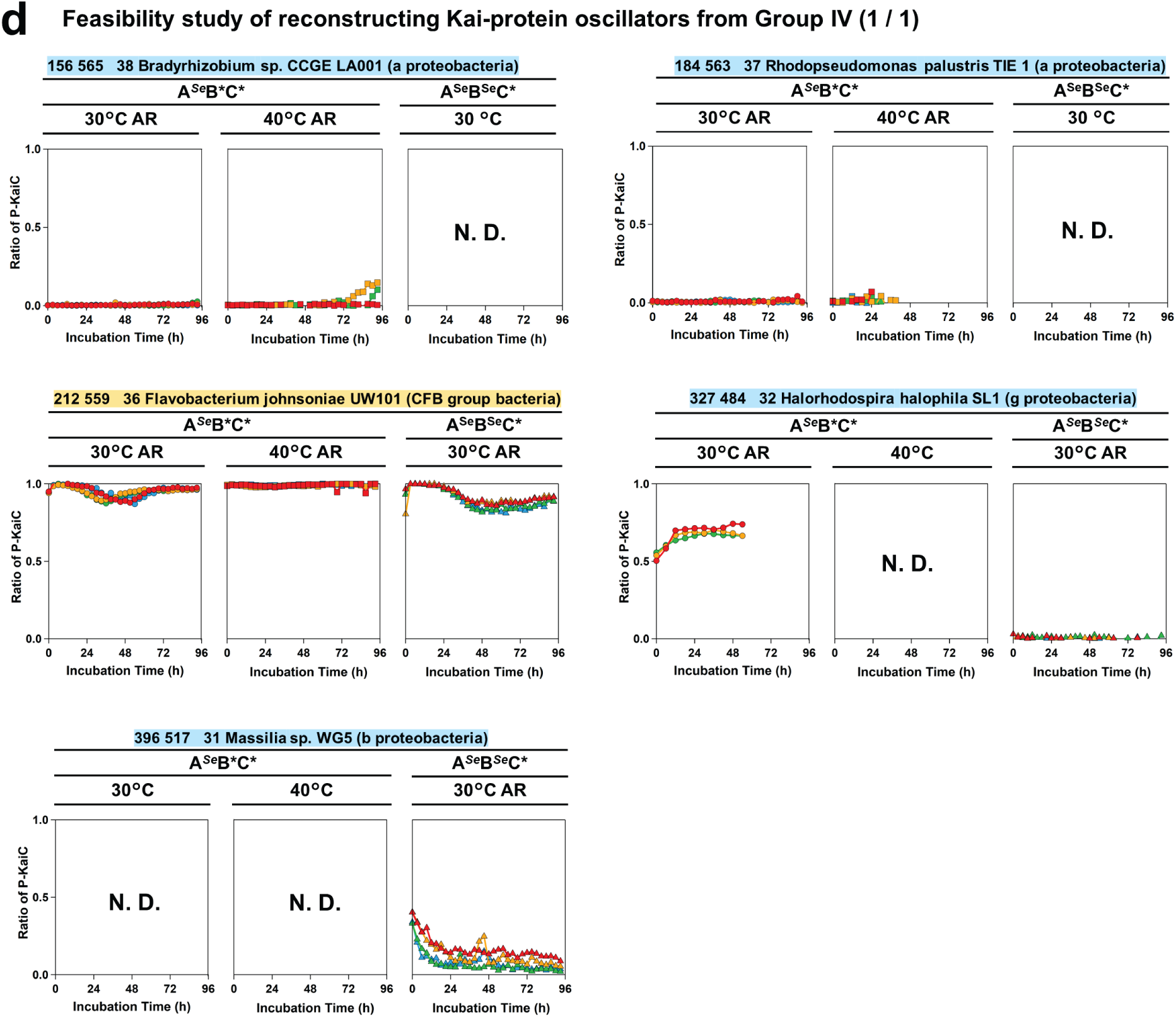
Feasibility study of reconstructing Kai protein oscillators in extant species. Each panel represents the results of *in vitro* rhythm assays for the A*B*C* or A*^Se^*B*C* set at 30°C (left) and 40°C (middle), and for the A*^Se^*B*^Se^*C* set at 30°C (right). Temporal profiles of the ratio of the phosphorylated KaiC (P-KaiC) were investigated using different KaiA concentrations 0.01 mg/mL (cyan), 0.02 mg/mL (green), 0.04 mg/mL (orange), and 0.08 mg/mL (red) but fixed concentrations of KaiC (0.2 mg/mL) and KaiB (0.04 mg/mL). The values listed near the measured temperature indicate the period length estimated from a temporal profile indicated by a symbol in parentheses. AR indicates that the period could not be determined (arrhythmic). N.D. indicates not tested.

**Extended Data Fig. 4.**
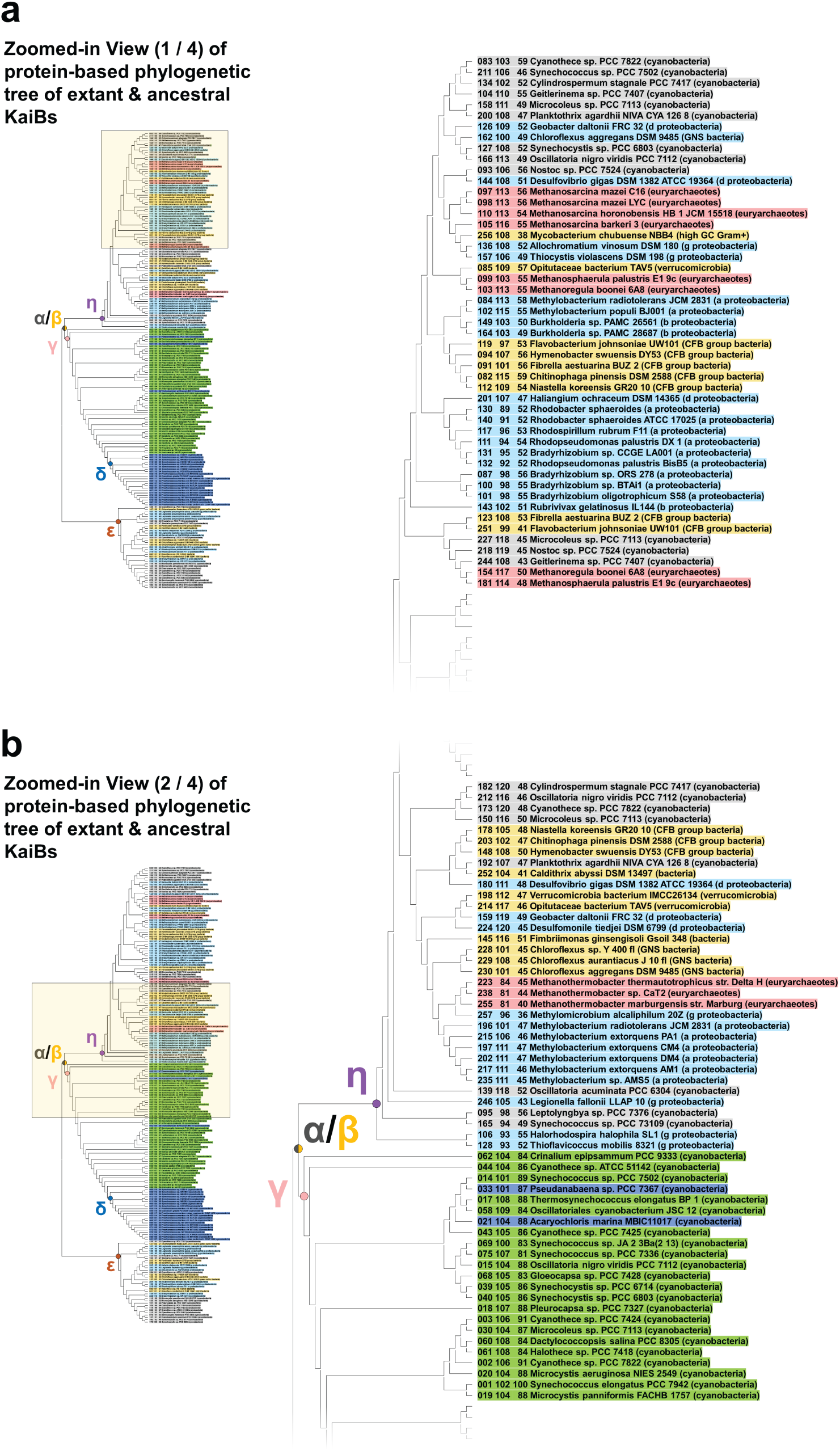
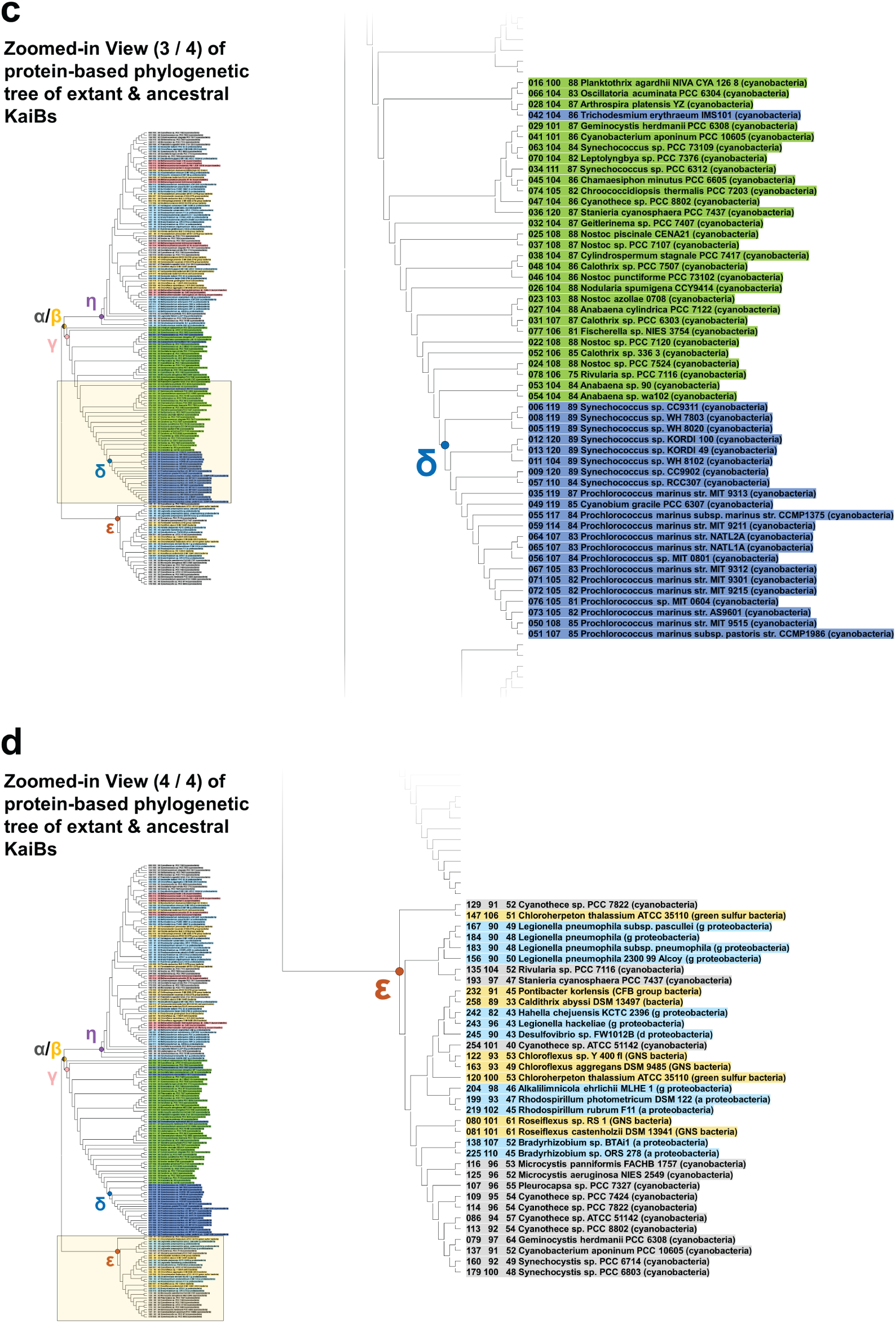
Phylogenetic tree of extant KaiB homologs. In each panel (**a**, **b**, **c**, and **d**), the enlarged image in the center corresponds to the area surrounded by a yellow square in the overall phylogenetic tree shown on the left. The three numbers preceding the name of each entry indicate the ID number, the total number of amino acids, and the sequence identity with KaiB*^Se^*, respectively. Cyanobacteria in freshwater, marine cyanobacteria, proteobacteria, bacteria, and euryarchaeota/crenarchaeota are colored green, blue, cyan, yellow, and red, respectively. Extant KaiBs from freshwater cyanobacteria (gray entries), which are located downstream of KaiB^ε^ and KaiB^η^, are originated from secondary copies or lateral transfer of *kaiB* genes downstream of KaiB^γ^, and thus can be excluded when identifying nodes based on species and growing habitant.

**Extended Data Fig. 5.**
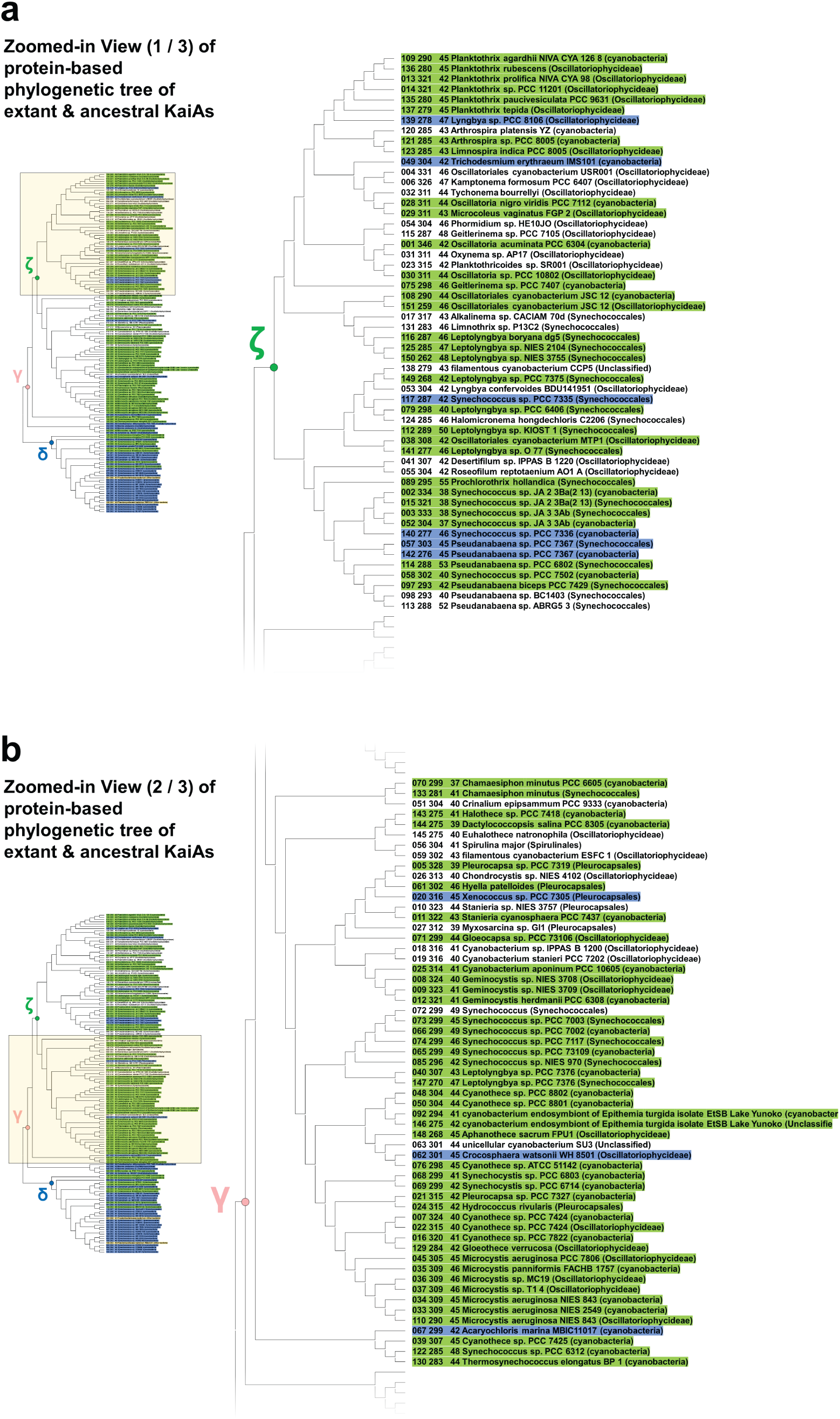
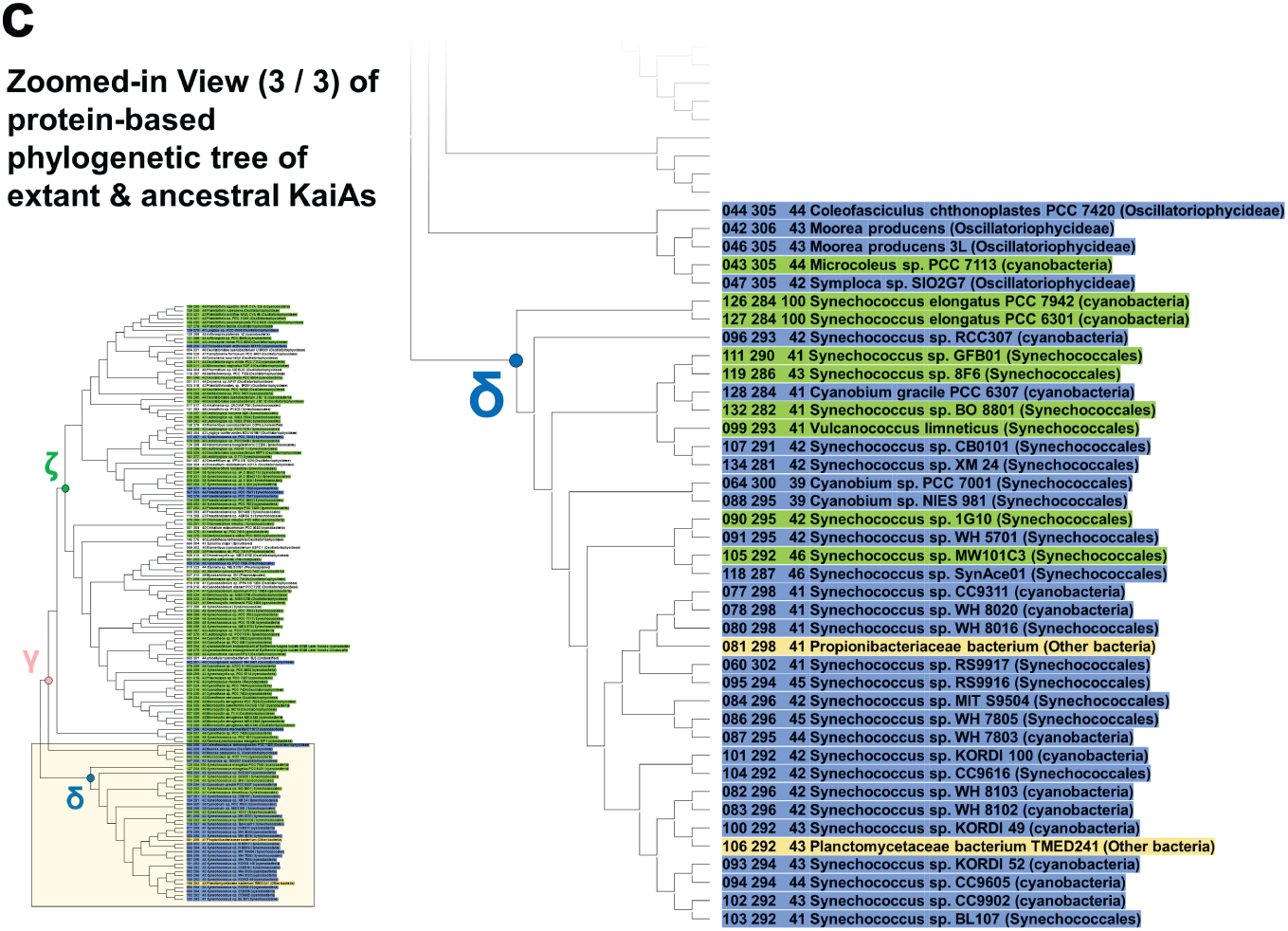
Phylogenetic tree of extant KaiA homologs. In each panel (**a**, **b**, and **c**), the enlarged image in the center corresponds to the area surrounded by a yellow square in the overall phylogenetic tree shown on the left. The three numbers preceding the name of each entry indicate the ID number, the total number of amino acids, and the sequence identity with KaiA*^Se^*, respectively. Cyanobacteria in freshwater, marine cyanobacteria, and bacteria are colored green, blue, and yellow, respectively.

**Extended Data Fig. 6.**
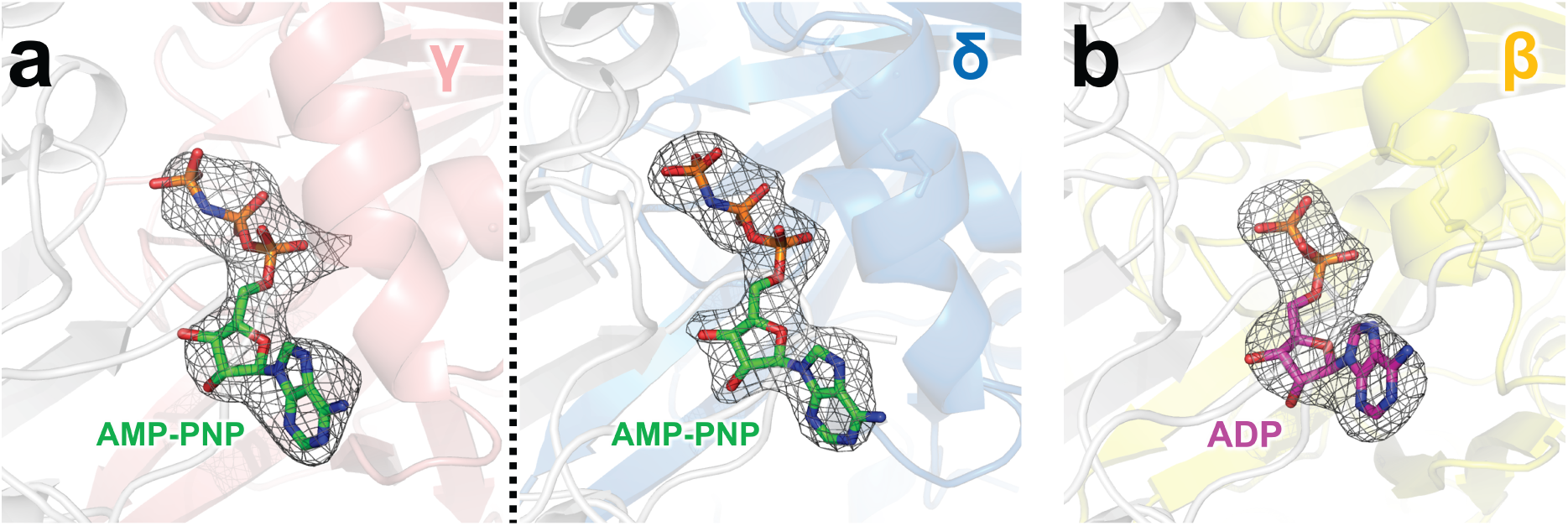
Nucleotides bound to the active site of the CI domains. (**a**) KaiC^γ^ (left) and KaiC^δ^ (right). (**b**) KaiC^β^. The meshes represent the *F*_obs_ − *F*_calc_ omit maps contoured at 3σ.

**Extended Data Table 1.**
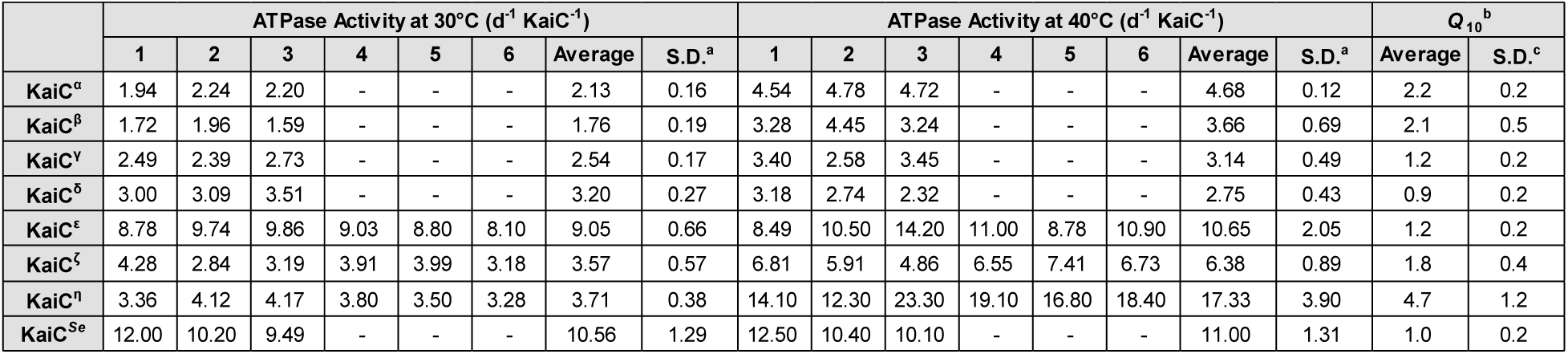
ADP production rates of ancestral KaiCs and KaiC*^Se^*. ^a^Standard deviations of the ATPase activity. ^b^*Q*_10_ values calculated as the ratio of the ATPase activities at 40°C to 30°C. ^c^Standard deviations of the *Q*_10_ values.

**Extended Data Table 2.** Data collection and refinement statistics. ^a^Values in parentheses correspond to the highest-resolution shell. ^b^*R*_merge_ = Σ|*I* − | / Σ*I*, where *I* corresponds to the observed intensity of reflections. ^c^*R*_work, free_ = Σ(|*F*_obs_| − |*F*_calc_|) / Σ|*F*_obs_|. *R*_free_ is the cross-validation of the *R*-factor using the test reflections, 5% of the data, not included in the refinements.

| Protein | KaiC<sup>β</sup> | KaiC<sup>γ</sup> | KaiC<sup>δ</sup> |
| --- | --- | --- | --- |
| Crystallization | 18mM Acetic acid,<br>65mM Sodium malonate | 0.94M Sodium formate,<br>1.20M Sodium malonate | 0.1M Tris-HCl (pH7),<br>1M KCl,<br>1.14M Sodium malonate |
| Cryo-protectant | 30mM Acetic acid,<br>95mM Sodium<br>malonate, 30% PEG4K |  |  |
| Nucleotide Addition | 5 mM AMP-PNP |  |  |
| X-ray Source | SPring-8 BL44XU |  |  |
| Data Collection |  |  |  |
| Space group | *P*2<sub>1</sub> | *P*1 | *P*2<sub>1</sub> |
| Unit cell length *a*, *b*, *c* (Å) | 93.1, 385.6, 108.1 | 90.8, 110.3, 166.9 | 92.0, 175.2, 94.8 |
| Unit cell angle α, β, γ (°) | 90.0, 113.2, 90.0 | 78.0, 87.3, 82.4 | 90.0, 96.9, 90.0 |
| Wavelength (Å) | 0.9 | 0.9 | 0.9 |
| Resolution range (Å)<sup>a</sup> | 49.30 - 3.00 (3.11-3.00) | 48.74 - 2.94 (3.05-2.94) | 49.19 - 2.54 (2.63-2.54) |
| Total reflections | 971459 (99836) | 868007 (50840) | 686983 (67782) |
| Unique reflections | 138997 (13900) | 132294 (12958) | 98128 (9618) |
| Redundancy | 7.0 (7.2) | 6.6 (3.9) | 7.0 (7.1) |
| Completeness (%) | 99.9 (100) | 99.5 (97.4) | 99.7 (97.9) |
| (I)/sigma(I) | 12.4 (2.1) | 12.3 (2.5) | 13.3 (2.6) |
| *R*<sub>merge</sub> (%)<sup>b</sup> | 0.12 (1.09) | 0.09 (0.51) | 0.09 (0.71) |
| Model building |  |  |  |
| Molecular replacement | 2GBL | 2GBL | 2GBL |
| Total atoms | 36725 | 41205 | 20931 |
| Protien | 36026 | 40387 | 20443 |
| Ligands | 672 | 764 | 384 |
| Water | 27 | 54 | 104 |
| *R*<sub>work</sub> (%)<sup>c</sup> | 28.0 | 23.9 | 22.5 |
| *R*<sub>free</sub> (%)<sup>c</sup> | 32.9 | 29.2 | 27.3 |
| R.M.S.D. from ideality |  |  |  |
| Bond length (Å) | 0.002 | 0.002 | 0.002 |
| Bond angles (°) | 0.61 | 0.64 | 0.77 |
| Average B factors (Å<sup>2</sup>) | 86.1 | 78.4 | 66.4 |
| Ramachandran plot |  |  |  |
| Most favored (%) | 91.2 | 95.0 | 96.7 |
| Allowed (%) | 7.2 | 4.6 | 3.2 |
| Disallowed (%) | 1.6 | 0.4 | 0.1 |
| PDB code | 8ZN5 | 8ZN6 | 8ZN7 |

**Extended Data Table 3.**
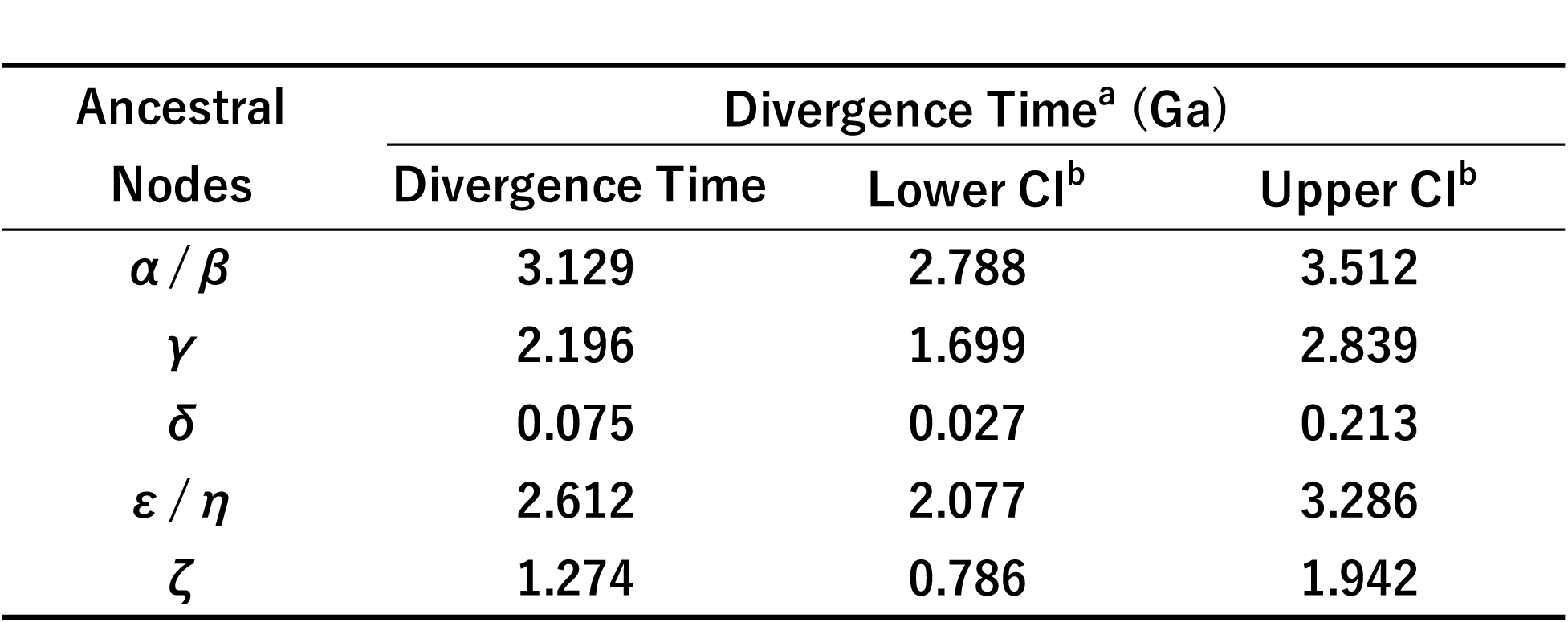
Divergence times of ancestral nodes in 16S rRNA–based phylogenic tree. ^a^TimeTree ^46^ estimate of the divergence times between *Synechococcus* and *Flavobacterium* was used to establish a calibration point at the node corresponding to the α/β point in the 16S rRNA–based phylogenic tree (Extended Data Fig. 2) as normal distribution with a median of 3.134 Ga and a deviation of 0.5 Ga. ^b^Confidence intervals.

