## Supplementary Information for "Evolutionary Origins of Self-sustained Kai Protein Circadian Oscillators"

#### Classification into groups I–IV and major entries in each group

To confirm the reliability of the classification of KaiC homologs into groups I–IV according to amino acid sequences, we constructed a 16S rRNA–based phylogenetic tree using 16S rRNA sequences from 108 entries of the 151 KaiC homologs used for the protein-based tree (Fig. 1a). The topology of the 16S rRNA–based tree (Extended Data Fig. 2) was similar to that of the protein-based tree of double-domain KaiC homologs (Fig. 1a and Extended Data Fig. 1); four major groups corresponding to cyanobacteria in freshwater, cyanobacteria in seawater, other bacteria, and other bacteria including archaea were reproducibly identified.

In the protein-based tree of the KaiC homologs (Fig. 1a and Extended Data Fig. 1), group I included 58 entries, including KaiC from *Synechococcus elongatus* PCC 7942 (KaiC$^{Se}$, ID: 001) [16]; KaiC from *Thermosynechococcus elongatus* BP-1 (KaiC$^{Th}$, ID: 025), which is often used in structural analyses [27]; KaiC from *Cyanothece sp*. ATCC 51142 (KaiC$^{Cy1}$, ID: 010), which has been studied in relation to ultradian rhythms [59]; and other KaiCs from major cyanobacterial genera such as *Microcystis*, *Synechocystis*, *Oscillatoria*, *Gloeocapsa*, *Nostoc*, and *Anabaena*.

Of 29 cyanobacteria in seawater, three (*Acaryochloris marina* MBIC11017 [ID: 066], *Trichodesmium erythraeum* IMS101 [ID: 006], and *Pseudanabaena sp.* PCC 7367 [ID: 076]) were exceptionally classified into group I, whereas the remaining 26 entries were classified into group II. Group II contained two genera, *Synechococcus* in seawater and *Prochlorococcus*. A previous study suggested that KaiC from *Prochlorococcus marinus subsp. pastoris str.* CCMP1986 (KaiC$^{Pr}$, ID: 080) functions as an hourglass-type oscillator with KaiB$^{Pr}$ in the absence of KaiA$^{Pr}$ [60].

Group III consisted of 13 entries from cyanobacteria and 10 entries from other bacteria. Some cyanobacteria possess multiple copies of *kaiC* genes [61]. Each of 13 cyanobacteria possesses multiple copies of *kaiC* genes, one of which is always the group I *kaiC* gene showing high sequence identity to the *kaiC* gene from *Synechococcus elongatus* PCC 7942. For example, *Cyanothece sp.* ATCC 51142 has a secondary copy as group III *kaiC* (KaiC$^{Cy2}$, ID: 085) in addition to group I *kaiC* (KaiC$^{Cy1}$, ID: 010). Comparison of the 16S rRNA–based and KaiC-based phylogenic trees suggest that 13 entries of *kaiC* gene from cyanobacteria listed in group III are the consequence of lateral gene transfer.

Group IV included three entries from archaea, six entries from cyanobacteria (*Geitlerinema sp.* PCC 7407 [ID: 126], *Cylindrospermum stagnale* PCC 7417 [ID: 147], *Cyanothece sp.* PCC

7822 [ID: 155], *Synechococcus sp.* PCC 7502 [ID: 154], *Oscillatoria nigro viridis* PCC 7112 [ID: 149], and *Synechocystis sp.* PCC 6803 [ID: 177]), and 33 entries from other bacteria. Each of these six cyanobacteria also has additional *kaiC* genes that are highly homologous to that from *Synechococcus elongatus* PCC 7942. Thus, the six entries of *kaiC* gene from cyanobacteria listed in group IV are possibly due to lateral gene transfer. KaiC from *Rhodobacter sphaeroides* ($KaiC^{Rs}$, ID: 194), which forms a dumbbell-like homo-dodecamer with $KaiB^{Rs}$ [34], and KaiC from *Rhodopseudomonas palustris* TIE1 KaiC ($KaiC^{Rp}$, ID: 184), which is not self-sustained but enhances fitness in a rhythmic environment [35], were included in this group.

Two entries from archaea, KaiC from *Pyrococcus horikoshii* OT3 (ID: 287) and KaiC from *Pyrodictium delaneyi* (ID: 309), were located in the outgroup (two bottom entries in Fig. 1a).

### pH and $Q_{10}$ values

The pH of the buffers listed in the main text and in Supplementary Information was measured at 25°C during buffer preparation. The p*K*a value of Tris is slightly temperature-dependent [62]. We confirmed that Tris buffers with pH 7.0 when prepared at 25°C change to pH 6.8 at 30°C and pH 6.6 at 40°C, and Tris buffers with pH 8.0 at 25°C change to pH 7.8 at 30°C and pH 7.6 at 40°C, as actual measured values at the corresponding temperatures. Therefore, $Q_{10}$ values calculated from data obtained at 30°C and 40°C using the corresponding Tris buffers could include additional pH-dependent effects, if any, of approximately –0.2 pH units.
